# Dosage compensation of transposable elements in mammals

**DOI:** 10.1101/2024.12.16.628797

**Authors:** Chunyao Wei, Barry Kesner, Uri Weissbein, Peera Wasserzug-Pash, Priyojit Das, Jeannie T. Lee

**Affiliations:** Department of Molecular Biology, Massachusetts General Hospital, Boston, MA, USA; Department of Genetics, Harvard Medical School, Boston, MA, USA

**Keywords:** Imprinted X inactivation, Random X inactivation, X hyperactivation, transposable elements, repetitive elements, noncoding, *Xist*, SINE, LINE, LTR, mouse preimplantation embryo, ES cell, escapee

## Abstract

In mammals, X-linked dosage compensation involves two processes: X-chromosome inactivation (XCI) to balance X chromosome dosage between males and females, and hyperactivation of the remaining X chromosome (Xa-hyperactivation) to achieve X-autosome balance in both sexes. Studies of both processes have largely focused on coding genes and have not accounted for transposable elements (TEs) which comprise 50% of the X-chromosome, despite TEs being suspected to have numerous epigenetic functions. This oversight is due in part to the technical challenge of capturing repeat RNAs, bioinformatically aligning them, and determining allelic origin. To overcome these challenges, here we develop a new bioinformatic pipeline tailored to repetitive elements with capability for allelic discrimination. We then apply the pipeline to our recent So-Smart-Seq analysis of single embryos to comprehensively interrogate whether X-linked TEs are subject to either XCI or Xa-hyperactivation. With regards to XCI, we observe significant differences in TE silencing in parentally driven “imprinted” XCI versus zygotically driven “random” XCI. Chromosomal positioning and genetic background impact TE silencing. We also find that SINEs may influence 3D organization during XCI. In contrast, TEs do not undergo Xa-hyperactivation. Thus, while coding genes are subject to both forms of dosage compensation, TEs participate only in Xi silencing. Evolutionary and functional implications are discussed.

## INTRODUCTION

In mammals, progressive gene loss on the Y chromosome during evolution resulted in dosage imbalance between X-linked genes in XY males and XX females, as well as a transcriptome-wide imbalance between X-chromosomes and autosomes^1–4^. To adapt to these inequities, two forms of dosage compensation have evolved. First, to balance X-chromosome dosages between males and females, female cells undergo X-chromosome inactivation (XCI) whereby one of the two X-chromosomes is transcriptionally silenced in females^5,6^. Because of XCI, males and females both have just one active X-chromosome (Xa). Thus, whereas all autosomes have two functioning homologues, the X chromosome has just one. This creates an X-to-autosome imbalance that necessitates the second form of dosage compensation: Hyperactivation of the active X-chromosome (Xa) leads to a near-doubling of Xa expression, to account for the relative X-linked deficit^7–10^ (Fig. 1a).

**Figure 1.**
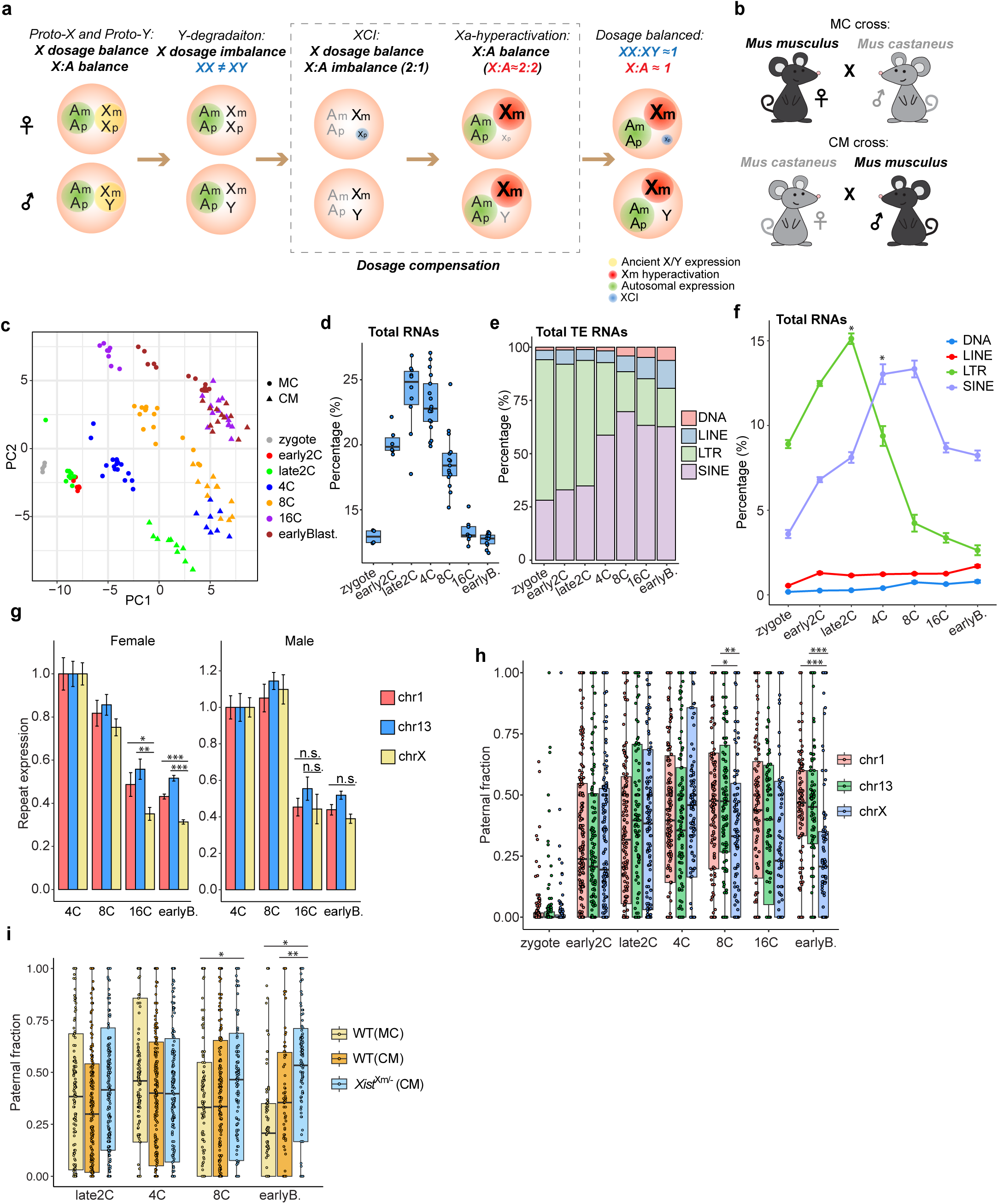
Dynamic TE expression in preimplantation embryos. a. A scheme describing mammalian dosage compensations consisting of XCI and Xa-hyperactivation. b. Two reciprocal interspecific crosses between *Mus musculus* (C57BL/6J) and *Mus castaneus* (CAST/EiJ) used in the study of preimplantation embryos. c. PCA analysis of wildtype preimplantation embryos of different stages from two reciprocal crosses. zygote and early 2C embryos were not collected in CM cross. d. Percentage of TE RNAs in the total transcriptome of MC cross preimplantation embryos. Total reads mapping to transposable elements (LINEs, SINEs, LTR and DNA) were compared with the total genome mapping reads (excluding rRNA reads). Centerlines in boxplots represent medians and box limits represent lower and upper quartiles. Whiskers represent 1.5x the interquartile range, and individual samples are represented as dots. Same box plot and dot plot representation are applied to all figures. e. Percentage of each TE class in the total expressed TE RNAs during preimplantation development (MC cross). f. Dynamic expression of 4 classes of TEs (LTR, SINE, LINE and DNA) in preimplantation embryos (MC cross). Percentage was calculated as relative to total transcriptome (depleted of rRNAs). \**p*<0.001, by paired t-test. g. Dynamic expression of autosomal TEs (chr1 and chr13) and X-linked TEs in female (left) and male (right) embryos. TE expression of each chromosome at each stage was normalized to 4C stage. Error bars represent s.e.m, n>=5 for each stage. \**p*=0.076, ** *p*=0.007, *** *p*<0.001, by paired t-test. *n.s.*, not significant. h. Silencing dynamics of TEs on the Xp in MC crossed embryos. Two autosomes (chr1 and chr13) are used as autosome comparators. Each dot on this graph represents a TE. \**p*=0.011, \*\**p*=0.002, \*\*\**p*<<0.001, by Mann-Whitney U test. i. The silencing dynamics of Xp TEs in wildtype embryos of reciprocal crosses versus female *Xist^Xm/-^* embryos . \**p*<<0.001, \*\**p*=0.008, by Mann-Whitney U test.

In mice, XCI occurs in two waves. It first takes place in preimplantation embryos, where the paternal X chromosome (Xp) is imprinted for silencing^11,12^. Most Xp genes are *de novo* silenced during cleavage stages of preimplantation female embryos. However, recent analysis showed that evolutionarily older genes demonstrate constitutive Xp silencing from the time of zygotic gene activation (ZGA) and can be traced backwards in developmental time to meiotic sex chromosome inactivation in the male germ line^13^. Thus, older Xp genes appear to be transmitted in a “pre-inactivated” state. In the blastocyst, imprinted XCI is continuously maintained in the trophectoderm and later in extraembryonic tissues. In contrast, epiblast cells of the embryo proper reactivate the Xp and undergo a zygotically driven “random” XCI, whereby one X chromosome is randomly selected for silencing during epiblast development ^14–16^. Both XCI forms require expression of *Xist*, a long noncoding RNA expressed from the future inactive X chromosome (Xi)^17–19^. For Xa-hyperactivation, although its timing and extent continue to be debated, recent studies are converging on the idea that Xa-hyperactivation is a plastic process linked to XCI^9,13^. For imprinted XCI in preimplantation embryos, genes on the maternal X (Xm) become hyperactivated as corresponding genes on the Xp become silenced. For random XCI in differentiating ES cells, Xa-hyperactivation is tightly linked to Xi silencing on a gene-by-gene basis. In male embryos, Xa-hyperactivation occurs constitutively at or shortly after ZGA^13^.

Although many transcriptomic studies have been carried out for XCI and Xa-hyperactivation^9,13,20–28^, conclusions have been limited to gene elements, which altogether constitute less than 2% of the X-chromosome^29^. On the other hand, non-genic elements account for the vast majority of the X-chromosome, of which TEs comprise half of its sequence. TEs are major components of genomic repeats. They are mobile genetic elements that occupy 40% of the mouse genome^30,31^. Based on the intermediate molecule used for mobilization, they are generally classified as DNA transposons or retrotransposons that rely on an RNA intermediate. While many have become evolutionary relics, some retrotransposons have remained active^32,33^. They insert into the host genomes through conversion of RNA intermediate to cDNA via reverse-transcription^34^. Retrotransposons can be further divided into two main categories — elements with long terminal repeats (LTRs), including retrovirus and LTR retrotransposons, and the elements without LTRs, such as long interspersed nuclear elements (LINEs) and short interspersed nuclear elements (SINEs). These repetitive elements are highly expressed and potentially contribute to a large fraction of transcriptome from the X-chromosome^35–37^.

Despite their high transcriptional content, TEs have not been profiled extensively in the context of XCI or Xa-hyperactivation. Part of the reason for the oversight has been the perception that TEs are parasitic elements with limited physiological function. There are also the technical challenge of capturing repeat RNAs, the difficulty of aligning repeats to the genome, and a dearth of analytical pipelines for determining allelic origin. However, emerging evidence indicates that TEs may have important genetic and epigenetic function. TE insertion or duplication is a major source of novel regulatory elements for the host genome^38^, such as promoters^31^, alternative splicing sites^39^ and insulator binding sites^30^. Furthermore, TEs have been associated with the stress response^40^. Their potential roles in regulating XCI have also been proposed, either at the initiation stage^41–43^ or in the spreading of *Xist* RNAs along the chromosome^44–47^.

Given this, here we set out to address the expression status of X-linked repetitive elements during dosage compensation. We had recently developed “So-Smart-seq” to overcome challenges associated with profiling a comprehensive transcriptome in the early embryo^13^. This method captures a more inclusive pool of RNA species, both poly-adenylated and non-polyadenylated; it reduces 5’ to 3’ coverage bias and is therefore advantageous for analyses that depend on SNPs, such as is the case for allelic analysis in XCI; and it preserves strand information and therefore accounts for both sense and antisense transcripts. Our analysis reveals unique silencing dynamics of X-linked TEs in both imprinted and random XCI, but finds that the TEs do not undergo Xa-hyperactivation. The evolutionary and functional implications are discussed.

## RESULTS

### Dynamic expression of TEs

Although many TEs (such as LINEs and SINEs) are believed to produce poly(A)-tailed RNAs, conventional single-cell RNA sequencing approaches have not yielded good TE coverages in preimplantation embryos (Supplementary Fig. 1a,b). We had previously used So-smart-seq to capture a more comprehensive transcriptome from preimplantation embryos and found extended coverage of both polyadenylated and non-polyadenylated RNAs^13^. The method previously enabled discovery of a class of X-linked genes that are inherited from the paternal germline in a pre-inactivated state ^13^. Here, we extended the analysis to transposable elements and asked whether they are also subject to dosage compensation. We profiled single preimplantation embryos at different stages to examine their gene expression pattern during imprinted XCI. To enable allele-specific analysis, we sequenced individual F1 preimplantation embryos at multiple stages including zygote (1-cell), early 2C (2-cell), late 2C (2-cell), 4C (4-cell), 8C (8-cell), and 16C (16-cell) and early blastocyst (earlyB., 32 to 64-cell) from interspecific crosses in both directions — (i) *M. musculus x M. castaneus* (*mus* x *cast*, MC) and (ii) *M. castaneus* x *M. musculus* (*cast* x *mus*, CM) (Fig. 1b). Using expression data from 1,200 different TEs, principal component analysis (PCA) and hierarchical clustering showed high similarity among replicates and clear separation of embryos by developmental stages. Surprisingly, embryos were also separated by crosses, suggesting a significant impact on TE expression by genetic background (Fig. 1c, Supplementary Fig. 1c). This is in sharp contrast with gene expression profiles, which are only affected by developmental stages^13^.

To study random XCI, we turned to mouse embryonic stem (ES) cells, as ES cells recapitulate random XCI during cell differentiation. Here we used a hybrid female mouse ES cell line that is *mus*/*cast* hybrid for chrX and chr13. This cell line selectively inactivates the X chromosome of *mus* origin due to a mutated *Tsix* on the *mus* X-chromosome^48^. Thus, the Xi is invariably of mus origin and the Xa of cas origin. To uncover TE expression during random XCI, we performed RNA-seq on differentiating ES cells from Day0 to Day10. Similarly, PCA using all expressed TEs showed sample separation consistent with the differentiation time (Supplementary Fig. 1d). These data indicate that our method sensitively captured unique TE expression patterns for each stage and cross, with high reproducibility among replicates.

In preimplantation embryos, our analysis showed significantly increased TE expression at the early 2C stage, coinciding with the minor wave of zygotic genome activation (ZGA) — much earlier than the expression of protein-coding genes, which largely occur during the major wave of ZGA at the late 2C stage. In both MC and CM crossed wildtype embryos, TE expression peaked at late2C and 4C stages, followed by a quick decline at later developmental stages, with the lowest expression level seen in early blastocyst embryos (Fig. 1d, Supplementary Fig. 2a). When total expression was stratified by TE class, we found dominant expression of LTRs from zygote to late2C stage, coinciding with the transition from maternal to zygotic expression in new embryos. Expression of LTR, however, quickly declined at the 4C stage and was supplanted by a wave of SINE expression (Fig. 1e,f, Supplementary Fig. 2b). These observations are in line with the reported dynamic change of global trimethylated lysine 9 on histone H3 (H3K9me3) and DNA methylation in early embryos^49–51^. In contrast, zygotic expression of LINEs and DNA transposons was very low (1∼2% of total transcriptome) in early embryos, with very little change during preimplantation development (Fig. 1f).

During random XCI in differentiating ES cells, we observed similar TE expression patterns. Total TE expression was first increased at the onset of cell differentiation and maintained at relatively high level until Day3, after which a continuous decline was observed. This is different from the stratified expression in preimplantation embryos. Relative expression from each TE class was stable throughout all stages during cell differentiation, with dominant expression from SINEs. Whereas LINEs were more abundant, DNA transposons remained low in expression in differentiating ES cells (Supplementary Fig. 2c,d). Altogether, our data reveal dynamic TE expression during preimplantation development and ES cell differentiation, with expression predominantly coming from LTRs and SINEs.

### Allelic determination of TE expression

To study dosage compensation of the Xi and Xa, we developed a bioinformatic pipeline to enable allelic expression analysis. Unlike genes, whose allelic expression can be determined by single nucleotide polymorphisms (SNPs) between two parental genomes, TEs are more challenging due to their multicopy nature and insufficient annotation of accurate SNPs and indels. Therefore, instead of relying on predefined SNPs or indels, we aligned sequencing reads to *de novo* assembled *cast* and *mus* genomes, and directly compared the alignment qualities of each read for *mus* versus *cast* reference genomes. To track the chromosome origin of TE-derived reads and to minimize bias, we only retained TE reads aligned uniquely to both *mus* and *cast*. Despite of the loss of many repeat-associated reads due to their multi-mapping status, we were able to retain allelic information for more than 3,900 TEs located across the entire X chromosome (Supplementary Fig. 2e). Using this pipeline, very few reads (if any) were assigned to the Xp in male embryos (Supplementary Fig. 2f), consistent with imprinted silencing of Xp. We verified our pipeline using bulk fibroblast RNA-seq datasets derived from pure *mus* and pure *cast* parental strains and observed exclusive mapping of *mus* and *cast* TE reads to their respective parental reference genomes (Supplementary Fig. 2g). Thus, allelic origin of TE-aligned reads can be determined using our pipeline with high confidence.

We then determined TE expression on autosomes versus X-chromosome in female preimplantation embryos. The expression of X-linked TEs mirrored that of autosomes until the 16C stage when chrX expression level became significantly lower than that of autosomes (Fig. 1g). This lower expression on the X, however, was not observed in male embryos. We therefore speculated that decreased TE expression on the X in the females may be related to Xp inactivation without compensatory Xa upregulation in females. To determine whether TEs were indeed subject to imprinted XCI, we calculated paternal fractions of all TEs on the X and autosomes in MC embryos. Consistently, most TEs were not paternally expressed until early2C stage. On X-chromosome, nearly all TEs were biallelically expressed at 4C, but showed progressively reduced Xp expression at the 8C stage and beyond. By the early blastocyst stage, TEs were predominantly repressed on the Xp (Fig. 1h). In contrast, autosomal TEs remained biallelic through preimplantation development (Fig. 1h, Supplementary Fig. 2h). In CM embryos, Xp silencing of TEs was initiated around the same stage as in MC embryos, but proceeded more slowly (Fig. 1i). This difference was not likely due to biased allelic analysis or comparison between inconsistent stages, as allelic dynamics of autosomal TEs were comparable in both MC and CM embryos (Fig. 1h, Supplementary Fig. 2i), and slow silencing of gene expression on the Xp was also observed in CM embryos (Supplementary Fig. 2j). Notably, Xp silencing of TEs was impaired in the paternal *Xist* knockout embryos, indicating an absolute requirement of *Xist* in TE silencing (Fig. 1i). We therefore conclude that TEs are subject to XCI during preimplantation development and require *Xist* for the initiation of silencing.

### SINE/LTR silencing and clustered escape on the Xp in preimplantation embryos

When further stratified based on class, X-linked TE RNAs were mainly expressed from three classes: SINEs, LTRs and LINEs (Supplementary Fig. 3a). Intriguingly, although all TEs were reactivated on the Xp around the 4C stage, they were different in timing and extent of silencing during imprinted XCI. Overall, LINEs and LTRs seemed to be sensitive to XCI in 8C embryos but showed strong heterogeneity throughout the rest of preimplantation development. In early blastocysts, LTRs were repressed to greater extent than LINEs. In contrast, all SINEs showed relatively similar silencing dynamics, exhibiting the strongest resistance to XCI among all three classes (Fig. 2a). Interesting, however, there was some paternal SINE expression from 1-cell zygotes prior to any ZGA (Supplementary Fig. 3b), in line with a recent study reporting that SINEs are rapidly demethylated in the paternal genome at the late pronucleus stage^52^.

**Figure 2.**
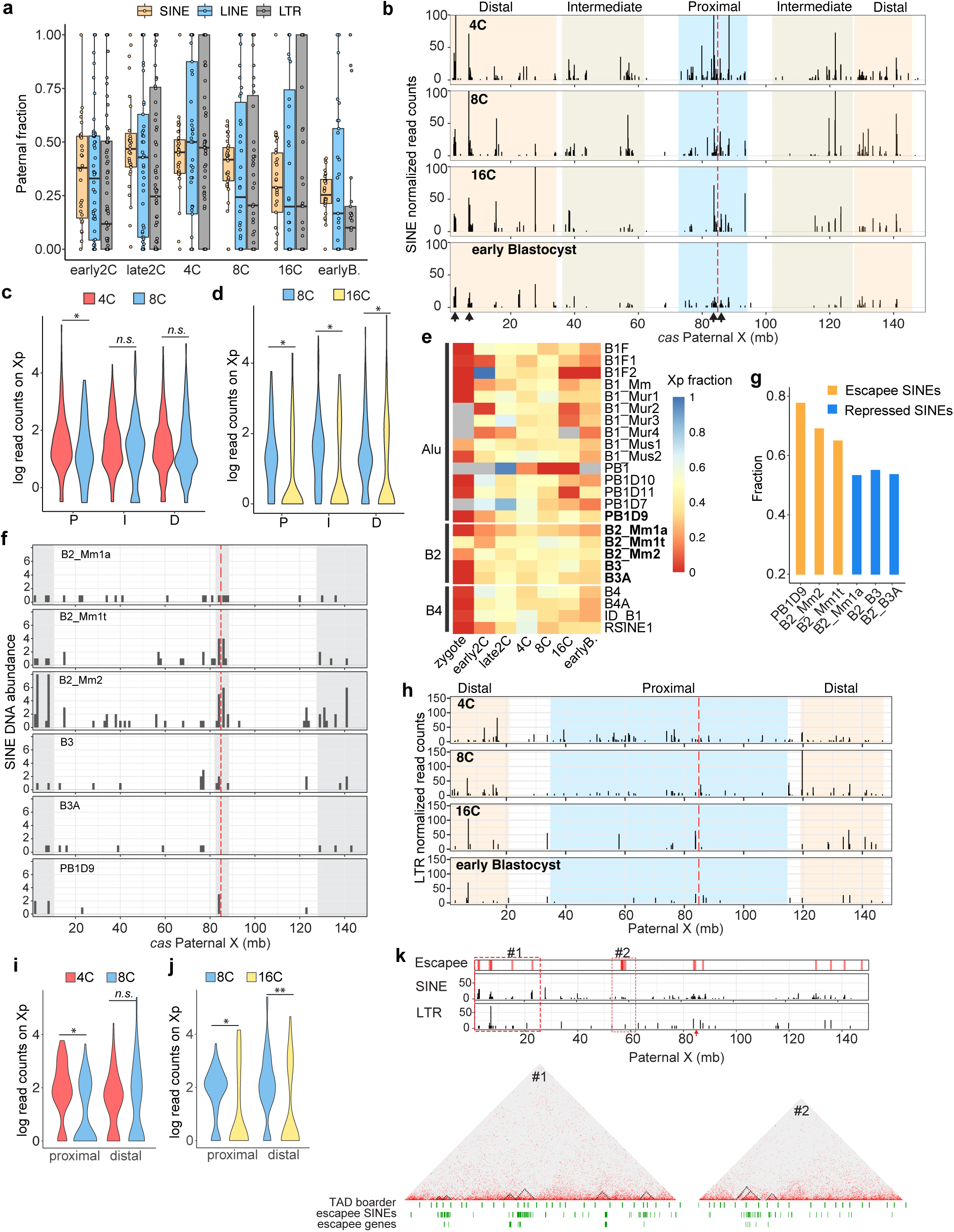
Silencing dynamics of Xp SINEs and LTRs in preimplantation embryos. a. Silencing dynamics of three classes of TEs (SINE, LINE and LTR) on Xp in preimplantation embryos. b. SINE expression declines across the entire Xp (*cas* allele) during imprinted XCI (from 4C to early blastocyst). The X-chromosome was divided into 3 regions (“Proximal”, “Intermediate” and “Distal”) by their proximity to *Xist* locus and labeled with different colors. Allelic read counts were normalized to the total expressed SINEs. The vertical red dash line marks *Xist* locus. Black arrows indicate the “hotspots” of SINE expression. c. Violin plots showing the Xp silencing of SINEs in different regions from 4C to 8C. P, Proximal; I, Intermediate and D, Distal. \**p*=0.0035, by Mann-Whitney U test. *n.s.*, not significant. d. Violin plots showing the Xp silencing of SINEs in different regions from 8C to 16C. Regions are the same as (C). \**p*<<0.001, by Mann-Whitney U test. e. Heatmap showing the paternal fraction of SINE expression on the X-chromosome during preimplantation development. Names of SINEs studied in the downstream analysis are marked in bold. f. Distribution of SINE loci marked in (E) across the entire Xp (*cas* allele). The vertical red dash line marks the *Xist* locus. Grey zones indicate the hotspots and active regions facilitating the escape of SINEs from imprinted XCI. g. Fractions of SINE DNA loci found in the hotspots and active regions on the Xp (grey zones shown in F). h. LTR expression across the entire Xp (*cas* allele) during imprinted XCI (from 4C to early blastocyst). X chromosome was stratified into 2 regions (“Proximal” and “Distal”) by their overall proximity to *Xist* locus and labeled with different colors. Allelic read counts were normalized to the total expressed LTRs. The vertical red dash line marks *Xist* locus. i. Violin plots showing the Xp silencing of LTRs in different regions from 4C to 8C. \**p*=8.64e-5, by Mann-Whitney U test. *n.s.*, not significant. j. Violin plots showing the Xp silencing of LTRs in different regions from 8C to 16C. \**p*=0.038, \*\**p*=0.0038, by Mann-Whitney U test. k. Bar graphs showing the overlap of escapee SINEs/LTRs with escapee genes in imprinted XCI. The red arrow marks *Xist* locus. Regions shown in the red boxes (#1 and #2) are further enlarged and revealed in the pseudo bulk Hi-C map obtained from the single-cell maps^55^.

To further study X-linked SINEs in preimplantation embryos, we collected all SINE allelic reads. Given that SINEs are often found in gene-rich regions, particularly gene introns, one concern would be that the SINE reads could be degradation products of host gene transcripts. However, metagene analysis showed that almost all SINE reads mapped to full SINE elements, with the 5’ ends precisely matching the SINE TSS (transcription start site; Supplementary Fig. 3c), arguing against their being derived from degradation of host transcripts. Because the allelic reads were uniquely aligned, we were able to map the reads to the X-chromosome and analyze dynamic changes in SINE expression along the entire X. As expected, SINEs on the Xp were progressively silenced through preimplantation development, whereas expression on the Xm remained unchanged (Fig. 2b, Supplementary Fig. 3d), consistent with imprinted XCI.

We found variable silencing dynamics among SINEs on the Xp. Interestingly, SINE silencing correlated with 2D proximity to the *Xist* locus. By linear distance to *Xist*, we divided the mapped SINEs into 3 categories: Proximal (within 15kb), Intermediate (15-57kb) and distal (further than 57kb) (Fig. 2b). Proximal SINEs initiated repression on the Xp as early as 8C stage, whereas farther SINEs did not until 16C stage (Fig. 2c,d). By the early blastocyst stage, SINEs in the proximal and Intermediate groups were all largely silenced, whereas distal SINEs still maintained expression, indicating that the timing and strength of SINE repression by imprinted XCI correlated strongly with linear proximity to *Xist.* Nevertheless, certain loci on the Xp appeared to be less affected by XCI in early blastocyst embryos, acting as potential “hotspots” for clustered SINE expression (Fig. 2b). Specifically, one such hotspot site was identified within the X inactivation center (Xic), a small region essential for initiating XCI in epiblast cells^53^.

We then stratified total SINEs by families using RepeatMasker. Overall, we found that most SINEs belonged to Alu, B2, and B4 families (Supplementary Fig. 3e). SINEs from different families, or even individual SINEs, showed distinct sensitivities to XCI in preimplantation embryos, with some B2 SINEs being more resistant to the Xp silencing (Fig. 2e). Of all B2 SINEs, two (*B2_Mm1t* and *B2_Mm2*) were least silenced on the Xp at early blastocyst stage, and *PB1D9* from Alu family was constitutively active (Fig. 2e). These elements therefore escape imprinted XCI. To investigate features distinguishing these SINEs from others, we located each B2 SINE on the Xp. Interestingly, escapee SINEs such as *B2_Mm1t* and *B2_Mm2* were not evenly distributed across the X-chromosome. Instead, they tend to cluster in “hotspot” regions, sometimes paradoxically clustering near the *Xist* locus (Fig. 2f, grey area). As *Xist* itself is an escapee, these locations may aid potentially in maintaining their active expression. In contrast, other more repressed B2 SINEs did not show such distribution and were much less enriched in these regions (Fig. 2f,g). This is also the case for escapee SINEs of other families, such as *PB1D9* in Alu family (Fig. 2f,g). Thus, SINE repression through XCI is highly affected by genomic locations.

We also observed a general association between the Xp silencing of LTRs and their linear distance to the *Xist* locus (Fig. 2h). LTRs located relatively close to the *Xist* locus on the Xp initiated repression at the 8C stage, whereas those further away from the *Xist* locus did not show signs of repression until the 16C stage or later (Fig. 2i,j). Silencing dynamics also varied among LTR families. While ERV1, ERVK, and ERVL LTRs were efficiently silenced, ERVL-MaLR LTRs were more resistant to XCI and emerged as major contributors to LTR expression on the Xp in early blastocyst embryos (Supplementary Fig. 3f).

Interestingly, when examining escapee SINEs and LTRs on the Xp in early blastocysts, we found that a large fraction of escapee elements resides in clusters, particularly around SINE “hotspots”. Furthermore, these clustered escapee SINEs and LTRs stayed in close proximity to identified escapee coding genes^13^ (Fig. 2k), implying a potential shared regulatory mechanism between these TEs and escapee genes. Considering the relationship between SINEs and 3D chromatin architecture^30,54^, we speculated that the clustering of active SINEs or LTRs on the Xp could lead to the formation of insulated regions that facilitate the sustained expression of escapee genes in a local repressive environment. To verify this hypothesis, we analyzed single-cell Hi-C data from early blastocyst embryos^55^ and found that clustered escapee SINEs tended to colocalize with defined topological domains on the Xp that also encompass escapee genes (Fig. 2k). Taken together, SINE and LTR elements on the Xp undergo progressive silencing through imprinted XCI, with their silencing dynamics strongly correlated with genomic locations and linear distance to *Xist*. Escapee SINEs tend to cluster on the Xp and colocalize in the same topological domains with escapee genes.

### Evolutionary age affects LINE silencing during imprinted XCI

Given the great heterogeneity in LINE silencing on Xp (Fig. 2a), we asked whether LINE silencing could be better dissected if further stratified. Our previous study had demonstrated that evolutionary age of genetic elements on the X could affect their silencing during XCI^13^. We thus refined our analysis accordingly and further categorized LINEs by evolutionary age into three major classes including (1) ancient mammalian LINEs (‘Old’) that emerged before mammalian radiation (L1M)^56^; (2) Murine LINEs (‘Mid’) that originated in the progenitor of modern murine rodents (Lx)^57^; and (3) mouse LINEs (‘Young’) that were found only in the house mouse (L1Md)^58^.

Indeed, silencing differed among the three LINE classes of different evolutionary age (Fig. 3a). L1M LINEs were very sensitive to XCI and showed strong suppression on Xp from 8C to early blastocyst stage. In sharp contrast, Lx LINEs were more resistant to XCI and did not show significant suppression even at the early blastocyst stage, although a general trend of Xp silencing was observed. Intriguingly, silencing of L1Md LINEs at the early blastocyst stage remained heterogenous, despite that they appeared to be efficiently silenced at 8C-16C stage (Fig. 3a). We then wondered how L1Md LINEs were expressed in the preimplantation embryos. In the mouse genome, full length LINEs are typically 5-6kb long and are found almost exclusively for L1Md LINEs on the X (Supplementary Fig. 3g), indicating that the L1M and Lx LINEs lost their independent transcription potential in the mouse. Indeed, most expressed L1M (Old) and Lx (Mid) LINEs recovered by our sequencing were less than 1kb in length (Fig. 3b), implying that they were mainly co-expressed with host genes such as pseudogenes due to truncated insertion. However, their expression was unlikely to depend on co-transcription with protein-coding genes, as both intronic and intergenic LINEs showed similar length distribution and silencing dynamics through preimplantation development (Supplementary Fig. 3h,i).

**Figure 3.**
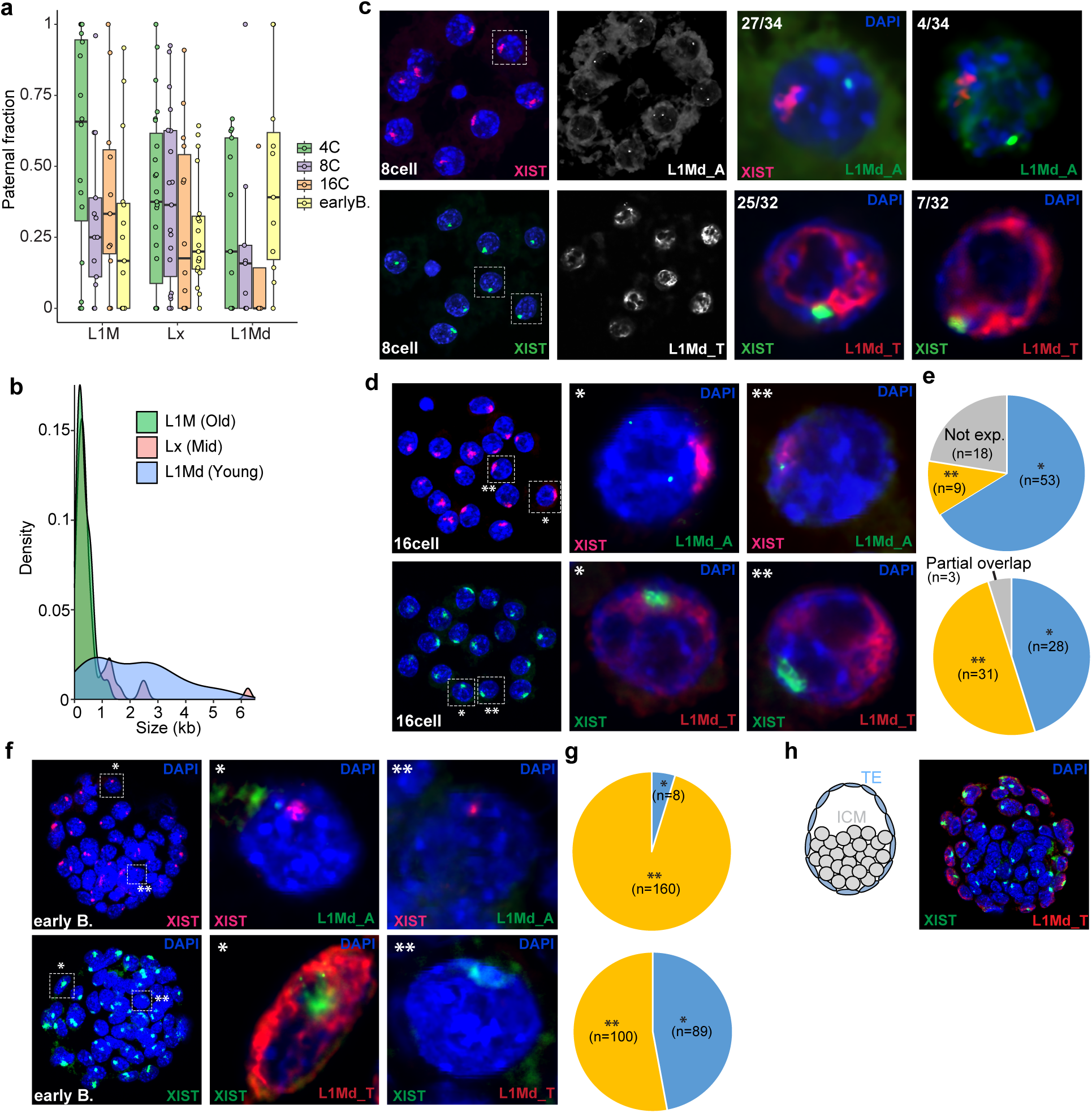
LINE silencing on Xp in preimplantation embryos. a. Silencing dynamics of 3 classes of LINEs with different evolutionary age. b. Length distribution of mapped LINEs in 3 classes. c. Nascent RNA FISH showing expression of two youngest LINEs (L1Md_A and L1Md_T) in 8cell embryos. *Xist* domain was labeled to indicate the location of paternal X. Representative cells squared in the left image are enlarged and shown to the right. d. Nascent RNA FISH showing the expression of two youngest LINEs (L1Md_A and L1Md_T) in 16cell embryos. *Xist* domain was labeled to indicate the location of paternal X. Two representative cells squared and marked by * or ** in the left image are enlarged and shown to the right. e. Quantification of observed embryos shown in (d) f. Nascent RNA FISH showing the expression of two youngest LINEs (L1Md_A and L1Md_T) in early blastocyst embryos. *Xist* domain was labeled to indicate the location of paternal X. Two representative cells squared and marked by * or ** in the left image are enlarged and shown to the right. g. Quantification of observed embryos shown in (f) h. (left) A schematic diagram describing a mouse preimplantation embryo at early blastocyst stage. (right) RNA FISH showing the spatial expression pattern of L1Md_T LINEs in early blastocyst embryos (n=4). *Xist* RNAs were labeled as a control.

In contrast to L1M and Lx LINEs, the mapped L1Md (Young) LINEs were much longer, with a large fraction ranging between 3-6kb in length (Fig. 3b), consistent with the enriched L1Md full-length LINEs on the X. This suggests that, unlike L1M and Lx LINEs that are mainly co-expressed with host genetic elements, L1Md LINEs were likely expressed via both co-transcription with host genes and independent transcription via their own promoters. However, since most of L1Md LINEs recovered in our data were also truncated to various extents, other approaches were needed to investigate the LINE expression from independent transcriptions.

We therefore designed probes specifically targeting the promoters of two youngest L1Md LINEs: L1Md_A (type A promoter) and L1Md_T (type T promoters)^59^, and performed RNA Fluorescent *in situ* hybridization (FISH) in 8C, 16C and early blastocyst female embryos. While L1Md_T was broadly expressed in cell nucleus, L1Md_A expression was relatively lower, and mostly restricted into few small areas, appearing like small puncta in the nucleus (Fig. 3c). By the 8C stage, the L1Md_A expression was already depleted on the Xp in most embryos while L1Md_T RNAs were only depleted in the *Xist* domain, with little to no overlap at the edge. In some rear cases, partial overlap of L1Md_T with *Xist* was observed (Fig. 3c). By the 16C stage, expression of L1Md_A was not observed on the Xp in all examined cells, suggesting a complete inactivation. For L1Md_T, expression was depleted on the Xp in half of the examined cells, while the rest cells still showed close contact of L1Md_T with *Xist* domain at the edge (Fig. 3d,e). In early Blastocyst embryos, expression of L1Md_T varied in different cells. While their RNAs were barely detected in the inner cell mass, expression remained robust in the trophectoderm (TE), where L1Md_T was still expressed at the outer rim of *Xist* domain, indicating the persistent expression at some regions of the Xp despite of the *Xist* mediated silencing (Fig. 3f-h). This differential LINE expression in these two cell lineages revealed by our FISH was not likely due to the difference in FISH probe penetration, as *Xist* probes clearly reached the cells in the inner cell mass (Fig. 3h). Thus, L1Md LINEs exhibited heterogenous silencing dynamics during imprinted XCI. While L1Md_A was rapidly silenced by the 16C stage, L1Md_T demonstrated a progressive, yet slow inactivation, with strong resistance even at early blastocyst stage. Altogether, our data suggest that dynamic silencing of LINEs in imprinted XCI is associated with their evolutionary age. The ancient L1M LINEs are silenced early during XCI, whereas murine Lx LINEs tend to remain active throughout the preimplantation development. Unexpectedly, the youngest L1Md LINEs on the Xp behave differently during XCI and in a cell type-dependent manner.

### Strain-specific TE silencing in imprinted XCI

Given the strong impact of genetic background on total TE expression in early embryos (Fig. 1c), we further sought to explore whether TE silencing in XCI had strain-specific affect. Analysis of each TE class in preimplantation CM embryos revealed an overall similar, yet much slower Xp silencing compared with MC embryos, particularly for SINEs (Fig. 4a,b). While the trend of slow Xp silencing was also observed for LINEs and LTRs, the difference between MC and CM embryos was not statistically significant, presumably due to the considerable heterogeneity of Xp silencing in these two repeat classes (Supplementary Fig. 4a,b). This delayed TE silencing on Xp in CM embryos was unlikely due to any potential bias in our analytical pipeline or uneven quality of *cast* and *mus* genomes, as similar delayed silencing also occurred in gene elements, which was determined using the annotated SNPs between two parental genomes and was independent of our analytical method (Supplementary Fig. 2j). Despite of the slow inactivation on the Xp, SINE silencing still differed significantly from that in *Xist* Knockout embryos with the same genetic background, which exhibited completely biallelic expression at early blastocyst stage (Fig. 4c, Supplementary Fig. 4c), whereas silencing of LINEs and LTRs in CM embryos was statistically indistinguishable from that observed in *Xist* knockout embryos, respectively (Supplementary Fig. 4d,e). When analyzing SINEs in a locus-specific manner, we found that the correlation still held true in CM embryos between SINE silencing dynamics and their linear distance to *Xist* locus. Similar to the analysis in MC embryos, we divided SINEs into proximal, intermediate and distal SINEs (Fig. 4d). From 4C to 8C stage, no significant change was observed for all SINE groups, although a slight decrease in read counts could be seen in proximal SINEs (Fig. 4e). From 8C to 16C, a significant decrease in SINE expression was visible in proximal and intermediate SINEs (Fig. 4f). By the early blastocyst stage, all SINEs started to show inactivation, with the majority of SINE expression remaining at the distal region of X chromosome, consistent with the Xp silencing in MC preimplantation embryos (Fig. 4d,g). This pattern also seemed to hold for LTR elements. LTRs relatively close to *Xist* tended to be silenced early during imprinted XCI, while those further away showed delayed silencing and were still expressed even at early blastocyst stage, the final stage for embryo-wide imprinted XCI (Supplementary Fig. 4f-h). Thus, our analyses reveal a consistent silencing pattern for SINEs and LTRs during imprinted XCI in both MC and CM embryos. Collectively, these data indicate that TE silencing in imprinted XCI occurs independently of genetic background, with a slower progression in CM embryos than in MC embryos.

**Figure 4.**
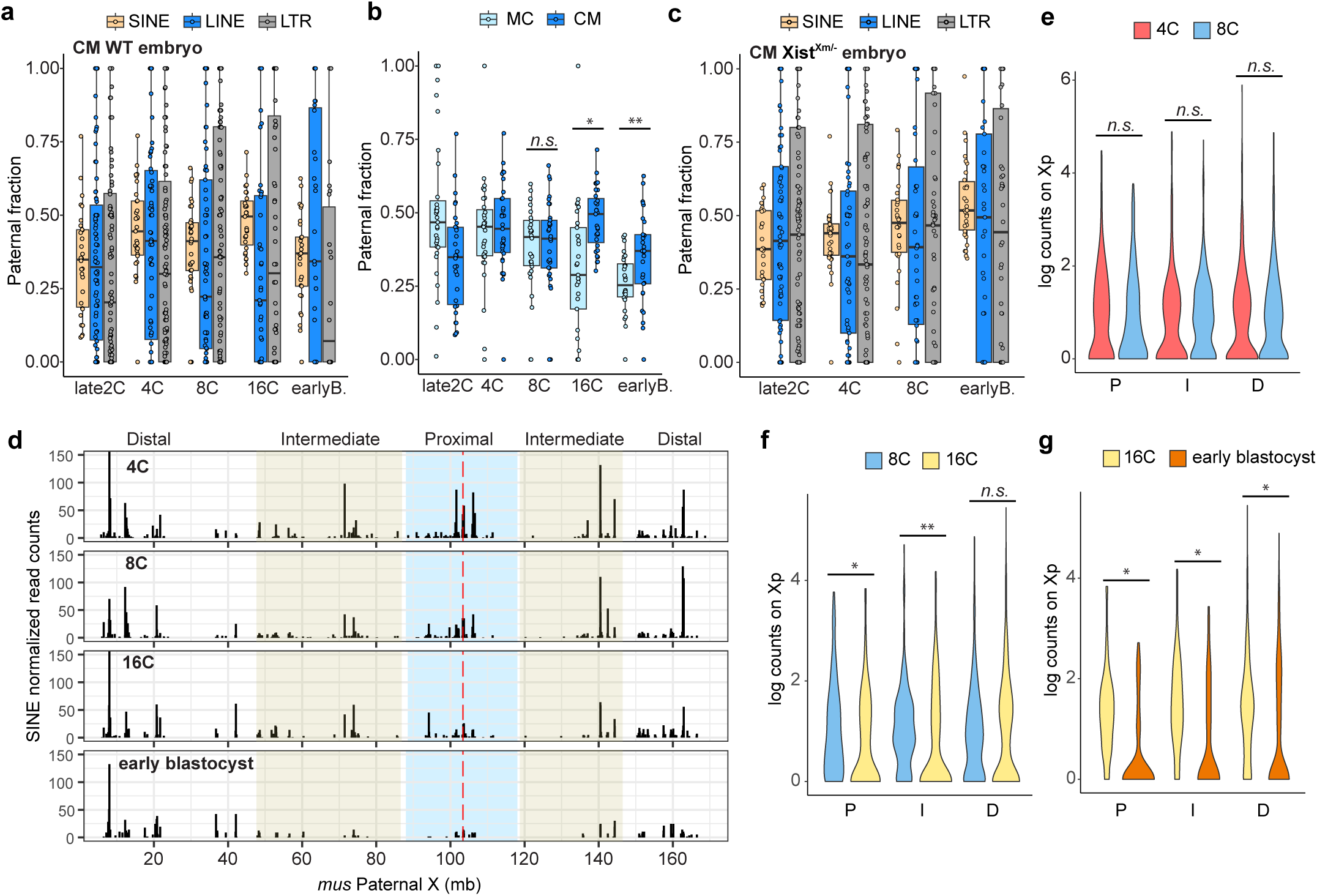
TE silencing in CM and paternal *Xist* knockout preimplantation embryos. a. Silencing dynamics of 3 classes of TEs (SINE, LINE and LTR) on Xp in CM wildtype preimplantation embryos. b. Silencing dynamics of Xp SINEs in both MC and CM wildtype preimplantation embryos. \**p*<0.001, \*\**p*= 5.49e-6, by Mann-Whitney U test. *n.s.*, not significant. c. Silencing dynamics of 3 classes of TEs (SINE, LINE and LTR) on Xp in paternal *Xist* knockout preimplantation embryos. d. SINE expression across the entire Xp (*mus* allele) in CM wildtype preimplantation embryos (from 4C to early blastocyst). X chromosome was stratified into “Proximal”, “Intermediate”, and “Distal” regions by the proximity to *Xist* locus, and labeled with different colors. Allelic read counts were normalized to the total expressed SINEs. The vertical red dash line marks *Xist* locus. e. Violin plots showing the Xp silencing of SINEs in different regions from 4C to 8C. *p* values were calculated by Mann-Whitney U test. P, Proximal; I, Intermediate and D, Distal. *n.s.*, not significant. f. Violin plots showing the Xp silencing of SINEs in different regions from 8C to 16C. \**p*= 0.003, \*\**p*= 7.68e-5, by Mann-Whitney U test. Regions are labeled the same as (E). *n.s.*, not significant. g. Violin plots showing the Xp silencing of SINEs in different regions from 16C to early blastocyst. \**p*<<0.001, by Mann-Whitney U test. Regions are labeled the same as (E)

### TE silencing is more complete during random XCI

We then sought to investigate whether TEs are silenced in in differentiating female ES cells, where random XCI is induced by cellular differentiation over 10 days. This female mouse ES cell line is *mus*/*cast* hybrid for chrX and chr13, with other chromosomes in pure *mus* genetic background. When considering all TEs combined, we found that the TE expression ratio on Xi (*mus* allele) remained relatively unchanged until Day2 of differentiation. Subsequently, TEs progressively lost their expression on Xi as differentiation proceeded, while chr13 remained constantly biallelic (Fig. 5a). By Day8, TE silencing was complete, as no further silencing was observed beyond this day. In contrast, other autosomes, as exemplified by chr1, were expressed entirely from the *mus* allele. In contrast to imprinted XCI, where silencing dynamics varied greatly among different TE classes, random XCI surprisingly revealed almost identical silencing dynamics on Xi for all TE classes, although SINEs were still slightly more homogenous than other TE classes (Fig. 5b). To further characterize TE silencing in random XCI, we mapped TE reads on the inactive (*mus*) X-chromosome and analyzed each class separately from Day0 to Day10 of differentiation. In SINEs, the silencing dynamics was very similar for almost all SINEs across the entire the X chromosome, regardless of their linear distance to the *Xist* locus (Supplementary Fig. 5a), implying distinct patterns of *Xist* spreading on the X between random XCI and imprinted XCI. In fact, majority of SINEs were already silenced by Day4. By Day8, SINE expression was completely shut down almost everywhere across the entire Xi. In line with this observation, SINE silencing dynamics was very similar when we stratified SINEs into different families (Fig. 5c, Supplementary Fig. 5b). Thus, SINEs were more efficiently and completely inactivated during random XCI. However, we still found six SINEs escaping silencing even at Day10, all of which overlapped with gene elements. Of the six escapee SINEs, three (*B3*, *PB1D9* and *B2_Mm2*) lied in the introns of escapee genes (*Ddx3x* and *Eif2s3x*, respectively), while the others (*MIRc*, *B1_Mus1*, and *ID_B1*) were colocalized with genes subject to XCI (*Clcn5*, *Xiap* and *Flna*, respectively) (Supplementary Fig. 5a,c).

**Figure 5.**
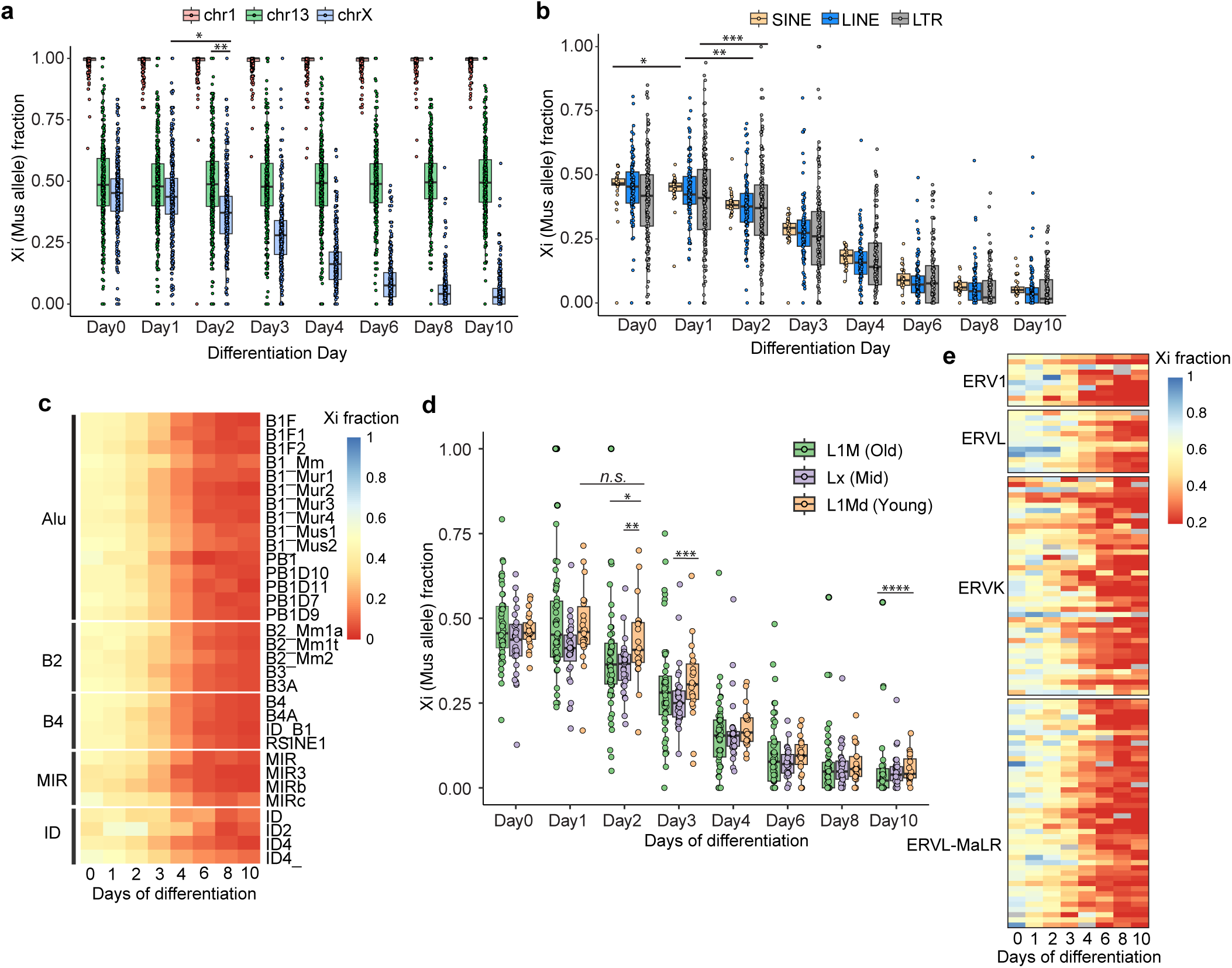
TE silencing in random XCI during ES cell differentiation. a. Silencing dynamics of TEs on Xi (*mus* allele) in differentiating ES cells. Two autosomes (chr1 and chr13) are used as autosome comparators. \**p*=0 4.97e-11, \*\**p*= 1.39e-23, by Mann-Whitney U test. b. Silencing dynamics of 3 classes of TEs (SINE, LINE and LTR) on Xi (*mus* allele) in differentiating ES cells. \**p*=0.029, \*\**p*= 5.40e-8, \*\*\**p*= 0.008, by Mann-Whitney U test. c. Heatmap of Xi (*mus* allele) fraction of 6 SINE families in random XCI during ES cell differentiation. d. Silencing dynamics of 3 classes of LINEs in random XCI during ES cell differentiation. \**p*=0.006, \*\**p*=0.005, \*\*\**p*=0.044, \*\*\*\**p*=0.007, by Mann-Whitney U test. *n.s.*, not significant. e. Heatmap of Xi (*mus* allele) fraction of 4 major LTR families in random XCI during ES cell differentiation.

For LINEs and LTRs, we observed similar silencing patterns in our chromosome wide analysis (Supplementary Fig. 5d,e). Almost all elements were silenced by Day 6, with a few escapees constantly being expressed at Day10 of differentiation. Most escapee TEs were colocalized with escapee genes, suggesting the establishment of a local transcriptional environment supporting TE expression (Supplementary Fig. 5c). When further stratified by evolutionary age into three major classes (L1M, Lx and L1Md), LINEs exhibited varied tendencies toward Xp silencing. Both L1M (Old) and Lx (Mid) LINEs started to show silencing at Day2, whereas L1Md (Young) LINEs did not initiate Xp silencing until Day3 (Fig. 5d). By Day 4, all three classes of LINEs progressed at the same pace toward Xp silencing. After 10 days of differentiation, L1M (Old) LINEs were significantly more repressed than L1Md (Young) LINEs, despite that inactivation had already been established for both LINE classes. Therefore, evolutionary age impacts LINE silencing in random XCI during ES cell differentiation. On the other hand, LTRs still exhibited varied silencing dynamics when stratified by families (Fig. 5e, Supplementary Fig. 5f). Taken together, these data show that TEs are subject to a more complete and thorough inactivation in random XCI than in imprinted XCI, and all TEs undergo silencing with similar dynamics. Additionally, evolutionary age affects LINE silencing in random XCI.

### Absence of Xa-hyperactivation for TEs

Lastly, we addressed the question of whether repetitive elements are subject to the second form of dosage compensation: Xa hyperactivation^4,7,8^. Recent study in mouse genes has demonstrated that Xa hyperactivation is closely linked to XCI and responds to Xi silencing on a gene-by-gene basis during both imprinted and random XCI ^9,13^. Given the active involvement of Xp silencing for TEs in two forms of XCI, we asked whether TEs were hyperactivated on the Xa for X:A dosage balance. Since total expression level of TEs on each chromosome was not available, instead of using X:A ratio to describe the status of Xa hyperactivation, we hypothesized that the presence of Xa hyperactivation would result in a larger TE expression increase (which was represented by the higher relative expression) on the Xa than that on autosomes, along with X inactivation.

To determine the chromosome-wide TE expression increase, we used the total normalized allelic read counts from each chromosome at each stage and calculated the relative expression by further normalizing to the beginning stage for the Xa and autosomes, respectively (see Methods). We first verified our hypothesis using gene elements, which has been known to undergo Xa hyperactivation upon Xi silencing. As expected, we found significantly higher relative gene expression on the Xa than that on autosomes starting at the 8C stage in preimplantation embryos, coinciding with the initiation of imprinted XCI (Supplementary Fig. 6a). Similarly, higher relative gene expression on the Xa was also observed in differentiating ES cells (Supplementary Fig. 6b). As a negative control, chr13 did not show significant difference when comparing to the rest autosomes (Supplementary Fig. 6c). Thus, gene analyses from both imprinted and random XCI validated our method.

Next, we analyzed relative expression of TEs. Surprisingly, in contrast to gene elements, which increased expression on the Xa upon Xi silencing, the relative expression of TEs was statistically indistinguishable between the Xa and autosomes at almost all stages in both male and female preimplantation embryos, indicating the absence of Xa hyperactivation for TEs during imprinted XCI (Fig. 6a,b). As expected, relative TE expression on chr1 was not statistically changed when comparing with the rest autosomes at almost all stages in female preimplantation embryos (Supplementary Fig. 6d). Therefore, while subject to Xp silencing in imprinted XCI, TEs lost Xa hyperactivation in preimplantation embryos. In differentiating ES cells, we found comparable TE relative expression on the Xa and autosomes until Day4 of differentiation, after which slightly higher relative TE expression seemed visible on the Xa than on autosomes, although not statistically significant (Fig. 6c). Interestingly, TE silencing on the Xi was almost completed by Day4 (Supplementary Fig. 5a,d,e). Furthermore, consistent with the chr1 analysis in preimplantation embryos, comparison between the chr13 and rest autosomes also revealed no statistical difference in relative TE expression during ES differentiation (Supplementary Fig. 6e). Thus, we conclude that the gene-by-gene based response of Xa hyperactivation to Xi silencing is absent for TEs, and that X:A dosage compensation is very weak, if at all, in differentiating ES cells. Altogether, our data reveal that cells are not sensitive to the change of X-linked TE dosage. In contrast to XCI, Xa hyperactivation does not occur for TE during either imprinted or random XCI.

**Figure 6.**
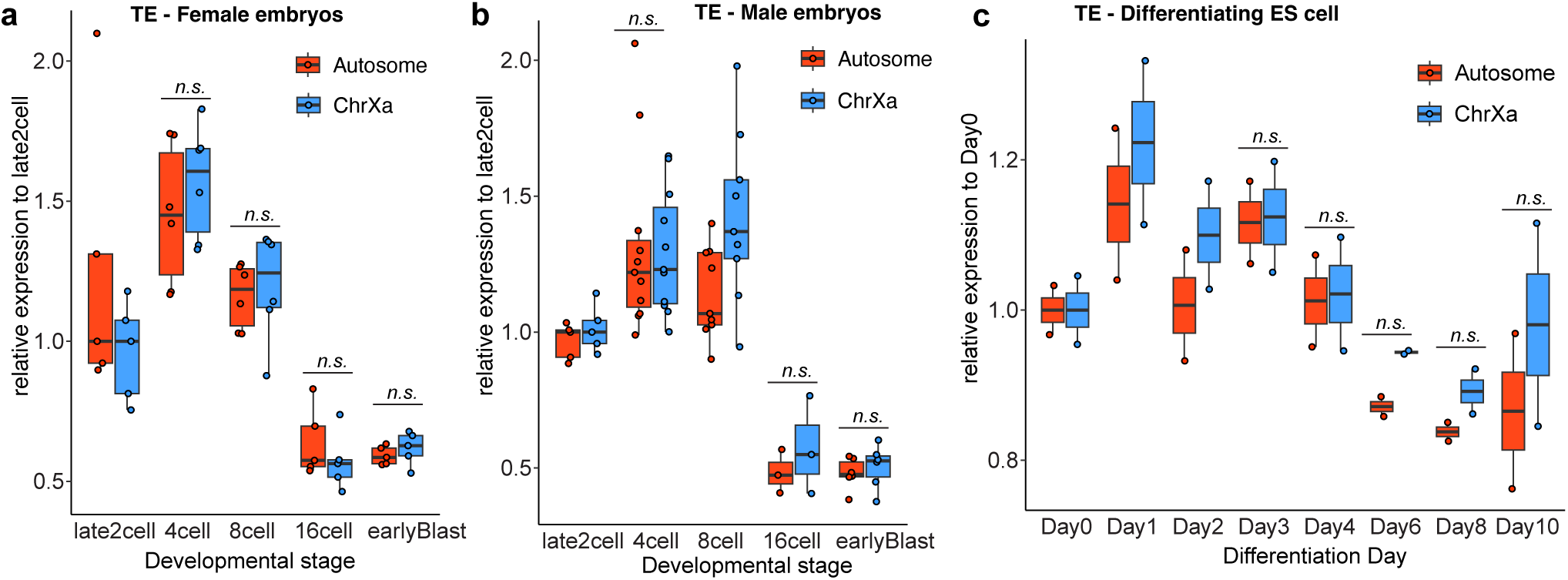
Xa hyperactivation is absent for TEs. a. The TE expression change (as represented by the relative expression comparing to late2C stage) on Xa versus autosomes during female preimplantation development. *n.s.*, not significant, by paired t-test. b. The TE expression change (as represented by the relative expression comparing to late2C stage) on Xa versus autosomes during male preimplantation development. *n.s.*, not significant, by paired t-test. c. The TE expression change (as represented by the relative expression comparing to Day0) on Xa versus autosomes during ES cell differentiation. P values were 0.126, 0.182 and 0.157 at Day6, Day8 and Day10, respectively, by paired t-test. *n.s.*, not significant.

## DISCUSSION

The mammalian X chromosome is low in gene density but rich in repeat elements, particularly TEs^29,60^. Because TEs can be highly transcribed, we argue that X-chromosome dosage should be measured across the whole chromosome inclusive of both genes and TEs. However, studies of X-linked dosage compensation, in either dynamic expression or mechanistic regulation, have largely overlooked TEs, despite their enrichment on the X. How they express and respond to XCI and Xa-hyperactivation has, until now, remained a significant gap in understanding. Here, we have examined these questions by developing a bioinformatic pipeline tailored to repetitive elements in an allele-specific manner. We then apply the new pipeline to So-Smart-Seq data generated from single embryos and ES cells for probing a comprehensive transcriptome and determining the allelic expression dynamics of three major TE classes on the X, including SINEs, LINEs and LTRs, during XCI and Xa-hyperactivation.

For both imprinted and random XCI, TE silencing absolutely requires *Xist*. We also note the influence of genetic background and chromosome loci on TE silencing dynamics. For imprinted XCI, the timing of Xp silencing on SINEs and LTRs is highly correlated with their linear distance to the *Xist* locus. Interestingly, for SINEs, the elements closest to *Xist* are the first to be silenced, potentially reflecting a closer proximity to the X-inactivation center in 3D space. However, for random XCI, TE silencing is more similar to gene silencing in that there does not appear to be a proximity effect, in line with the findings that *Xist* is spatially distributed and reaches multiple regions on the X simultaneously during the spreading^61,62^. Our study also uncovers TEs that escape XCI. During imprinted XCI, escapee SINEs/LTRs (or primarily SINEs) are frequently enriched in the regions adjacent to escapee genes, although they are not located within the genes. In random XCI, a few escapee SINEs are identified within introns of escapee genes. Two explanations can be offered for this association. The close linkage could reflect co-regulation by a 3^rd^ party element that protects the grouping of genes and SINEs from XCI. Alternatively, SINEs could potentially function as cis regulatory elements themselves (such as enhancers or insulator binding sites) for gene escape, perhaps through 3D architectural effects. Our findings align with previous studies reporting that SINEs are the dominant TEs contributing to chromatin accessibility and functional CTCF sites (including loop anchors) in the mouse genome^30,31^. Moreover, the RNA polymerase III (Pol III) promoter that drives SINE expression may also play roles in maintaining gene expression during XCI. In line with this speculation, one previous study indicated that Pol III transcription is not susceptible to *Xist*-mediated silencing, and inhibition of Pol III transcription affects gene reactivation in *Xist*-depleted fibroblast cells^63^. In the future, it would be interesting to investigate the timing and extent of Pol III depletion during XCI, as well as the roles of SINEs in constructing 3D genome structures.

Finally, our study demonstrates that, in sharp contrast to gene elements, TEs do not undergo Xa-hyperactivation. During preimplantation development, we observe no indication of Xa hyperactivation in both male and female embryos. In differentiating ES cells, Xa hyperactivation is clearly absent in the first 4 days of differentiation, by which TE silencing has almost completed on the entire Xp. This is in line with the absence of Xa hyperactivation noted in preimplantation embryos. Thus, with regards to X-to-autosome balancing, TE dosage inequality appears to be less consequential. During genome evolution, TEs provide major sources for genetic innovation, but also threaten genome stability^64^. Thus, a precise control of TE expression is important to ensure a delicate balance between expression and repression of TEs^65^. In preimplantation embryos or ES cells, the permissive TE expression due to global epigenetic reprogramming may result in ectopic TE dosage, which might reflect a “stress” state for cells. Thus, beyond the role of dosage balance between sexes, XCI may evolve as an additional important regulatory mechanism to repress overall TE dosage in cells. This may explain why XCI of TEs persists during evolution and early mouse development, but becomes uncouples from hyperactivation of any kind on Xa. Our observations suggest that, on the Xa, TEs are already active and may not need to be hyperactivated. However, we note that we did not examine Xa hyperactivation of TEs under induced stress. Under heat shock, for example, TEs such as SINEs become massively upregulated and play a crucial role during the stress response^40^. It is possible that some TEs would become hyperactivated on the Xa under such conditions. It is also possible that, unlike genes, the multicopy and ubiquitous nature of TEs may lead to less dependence on X-linked expression. The decreased TE expression on the X due to XCI could potentially be compensated by the enhanced expression of autosomal TEs. In summary, our study comprehensively determines the dynamic allelic expression of TEs in the context of two forms of X-linked dosage compensation and reveals a divergence in TE silencing between them. Our new bioinformatic pipeline and the resulting findings will provide insight for future TE research for X-linked dosage compensation and other epigenetic and allelic phenomena.

## ACKNOWLEDGEMENTS

We thank Carlos Rivera, Niklas Grimm and Roy Blum for critical reading of the manuscript, and all lab members for support. This work was funded by grants to J.T.L. from the NIH (R01-GM58839).

Imaging was performed in the Microscopy Core of the Program in Membrane Biology, which is partially supported by a Centre for the Study of Inflammatory Bowel Disease Grant DK043351 and a Boston Area Diabetes and Endocrinology Research Center (BADERC) Award DK135043. The AXR confocal imaging system is supported by NIH grant S10 OD032211-01.

## AUTHOR CONTRIBUTIONS

C.W. and J.T.L. conceived the project, analyzed data, and wrote the paper. C.W. performed all experiments and bioinformatics analyses. B.K. performed some bioinformatic analyses. C.W. and U.W. performed LINE RNA FISH experiments in preimplantation embryos. P.W. performed FISH imaging using confocal microscope, and P.D. performed the single-cell Hi-C analysis in embryos.

## DATA AVAILABILITY

The differentiating mouse ES cell RNA-seq data as well as processed TE expression data in preimplantation embryos and differentiating ES cells were deposited into GEO with accession number GSE275192. The single embryo So-Smart-seq data, including wildtype CM and MC embryos and paternal *Xist* Knockout embryos, are under GEO accession number GSE168455. The single-cell Hi-C data are under the GEO accession number GSE82185.

## METHODS

### Mice

All mouse experiments presented in this study were conducted in accordance with the animal research guidelines of NIH and approved by the Institutional Animal Care and Use Committee of Massachusetts General Hospital. All wild type preimplantation embryos were derived from reciprocal natural crosses between *C57BL/6J* and *CAST/EiJ*. The paternal *Xist* knockout embryos were obtained by mating wildtype *CAST/EiJ* females with *Xist*^-/Y^ males^18^ (*129S1/SvlmJ*). All used mice were at the age of between 8 and 12 weeks.

### Cell lines

The mouse ES cell line is a Mus (*129S1/SvlmJ*)/Cast *(CAST/EiJ)* F2 hybrid cell line (for chr13 and chrX) carrying a mutated *<u>Tsix</u>* allele that was previously described as “*Tsix*^TST/+^”^48^. ES cells were grown in feeder-free 2i medium (50% DMEM/F12 media, 50% Neurobasal media, 2% Hyclone FBS (Sigma), 0.5x N2 and 0.5x B27 supplements, 0.25x Glutamax, 100U/ml Pen/Strep, 0.1 mM βME, and 1000 U/mL ESGRO recombinant mouse Leukemia Inhibitory Factor (LIF) protein (Sigma, ESG1107)) supplemented with 1uM PD0325901 and 3uM of CHIR99021 (Selleck Chemicals) at 37°C with 5% CO_2_. ES cell differentiation was induced by replacing 2i medium with differentiation medium (DMEM, 10% FBS, 25mM HEPES, 1x MEM NEAA, 100U/ml Pen/Strep, 0.1 mM βME).

### Preimplantation embryo preparation

Embryos at the stage of 8cell, 16cell and early blastocyst (32-64 cell) were harvested at approximately E2.25, E2.75, E3.5, respectively. Each embryo was closely examined before experiments to ensure its normal morphology and correct number of blastomeres for the stage of interest. Embryo collections were performed by flushing oviduct and uteri with M2 medium (EMD Millipore) with a syringe and embryos were washed twice in M2 medium. To remove zona pellucida, each embryo was briefly incubated in Acid Tyrode’s solution (Sigma-Aldrich), followed by three washes in PBS containing 1mg/ml acetylated BSA (Sigma).

### mESC RNA-seq library preparation

Total cell RNA was extracted using TRIzol Reagent (Thermo Fisher Scientific), from which rRNAs were depleted using RiboMinus Eukaryote Kit v2 (Thermo Fisher Scientific) following manufacturer’s protocol. RNA-seq libraries were prepared in two biological replicates using NEBNext Ultra II Directional RNA Second Strand Synthesis Module and NEBNext Ultra II DNA Library Prep Kit for Illumina (New England BioLabs) as per manufacturer’s instructions.

### Library sequencing

All libraries were pair-end sequenced on the platform of Illumina Novaseq, following the manufacture instructions.

### So-smart-seq data analysis

Raw data was first examined using Trim Galore (The Babraham Bioinformatics Group) to remove adapters from reads using the following parameters: (--stringency 8 --phred33 -e 0.2 --paired --length 43--r1 44 --r2 44), followed by the PCR duplication removal based on sequences of attached RNA identifier and subsequent base trimming from the 5’end of reads (22nt from read1 and 5nt from read2). For gene analysis, VCF files reporting SNP sites of *CAST/EiJ* (Cast) based on mm10 were downloaded from Sanger Institute and used to reconstruct the Cast genomes from the mm10 genome assembly. This reconstructed Cast genome along with mm10 *C57BL/6J* mouse genome were used as parental genomes for read alignment. Pre-processed reads were aligned to two parental mouse genomes above separately using STAR (v2.7.10a) in the 2-pass mode, with the allowance of 1 mismatch in every 20 bases and a maximum of 6 mismatches per read pair^66^. Non-canonical introns were not considered, and read alignment spanning introns more than 60kb were filtered out. Reads with overlapping mates were only considered with at least 5 nucleotide overlap. All discordant read pairs where two mates were aligned to different chromosomes were discarded. In addition, two mates with more than 60kb gap in the alignment was ignored. Reads with more than 20 hits in the genome were ignored in the downstream analysis. The coordinates of rRNA genes in mm10 mouse genome were downloaded from RepeatMasker^67^ track (v4.0.7) in UCSC genome browser and all the reads mapping to these regions were masked and ignored in the final output files (including bam, bed and bw files). To quantify overall expression of each given gene, all uniquely aligned read pairs overlapping with gene exons were counted using featureCounts (v1.5.0-p1)^68^. For allelic analyses, each uniquely aligned read that covered strain specific SNPs or indels was analyzed and the alignment quality scores of each given read between two parental genomes were compared to determine its allelic origin. Pairs that differed significantly in alignment score due to mismatches/gaps were classified as allele-specific and the better alignment retained. Pairs with identical alignment scores or scores that differed only slightly due to fragment length penalties were classified as neutral. Each experiment yielded three tracks: paternal, maternal, and composite (neutral, paternal and maternal combined). For better visualization of gene expression between alleles, read mapping coordinates in different parental genomes were also converted to match coordinates in mm10 mouse genome for view in IGV^69^. Similar to overall expression quantification, only the allelic read pairs overlapping with gene exons were considered for allelic read counts of this gene.

For TE analysis, a *de novo* assembled *CAST/EiJ* mouse genome (GenBank assembly: GCA_921999005.2, Wellcome Sanger Institute) and mm10 *C57BL/6J* mouse genome was used as parental genome for read alignment. The read alignment via STAR was the same as gene analysis above, except that only reads with less than 200 hits in the genome were considered for the downstream analysis. The TE in the genomes (LINE, SINEs, LTR and DNA repeats) were determined by RepeatMasker^67^ with default parameters. For total repeat expression analysis, reads with alignment overlapping with annotated TE were retained and used to quantification. For allelic reads in TE, reads that did not overlap with any portion of the TE in either of the two gnomes were filtered out. Only the reads that were uniquely aligned in both two genomes and that overlapped with TE were considered for downstream analyses. Reads that were uniquely aligned in one genome but had multiple or no hits in the other gnome were also discarded. The alignment qualities of each single read between to parental genomes were compared to determine its allelic origin. The significance of mono allelic or biallelic expression of a given repeat element was determined using the Binomial distribution with Bernoulli *p*=0.5.

### mESC RNA-seq analysis

Adapters were removed from raw data using Trim Galore with the following parameters: (--stringency 5 -e 0.2 --length 35 --r1 36 --r2 36). For gene analysis, The VCF files reporting SNP sites of two parental strains *CAST/EiJ* (*Cast*) and *129S1/SvlmJ* (*mus*) based on mm10 were downloaded from Sanger Institute and used to reconstruct the Cast and mus genomes, respectively, from the mm10 genome assembly. For TE analysis, a *de novo* assembled *CAST/EiJ* genome (GenBank assembly: GCA_921999005.2) and *129S1/SvImJ* genome (GenBank assembly: GCA_921998555.2) were used as parental genomes for read alignment. After PCR duplicates removal, pre-processed reads were aligned to two parental mouse genomes separately using STAR (v2.7.10a) in the 2-pass mode, with the same parameters as used in the So-smart-seq alignment, except that a maximum of 8 mismatches were allowed per read pair.

### Allelic skewing analyses

Paternal expression fraction was used to represent the allelic skewing state for both genes and TEs on the chrX and autosomes. To increase the accuracy and credibility, genes having less than 13 total allelic reads in all replicates were ignored in the mESC RNA-seq data. At each stage, the paternal expression fraction of a given gene was calculated by averaging the paternal allelic fractions of this gene from two replicates. To perform allelic skewing analysis for TE on each chromosome, allelic reads mapping to a given TE (regardless of copy numbers) on one chromosome were summarized as the total allelic reads for this given TE on this chromosome. Paternal expression fraction of a given TE on a given pair of chromosomes was calculated as the ratio of total paternal allelic reads in relative to total allelic reads of this given TE on the given pair of chromosomes, similar to gene allelic analysis. TEs with less than 5 total allelic reads on one pair of chromosomes were filtered out from this chromosome, and TEs that were detected on less than five autosomes or TEs that were not detected on X chromosomes in all replicates were also discarded. TE allelic ratio was then calculated by averaging the allelic ratio of each given TE from all qualified replicates. For skewing density analysis of Old, Mid and Young LINEs, allelic reads in all replicates at each stage were combined to increase the allelic read number. All the boxplots and density plots were generated in R (4.0.2) using ggplot2^70^ (v3.3.2)., and all heatmaps were generated in R using pheatmap (v1.0.12).

### Single-cell Hi-C analysis

To construct the bulk allele-specific Hi-C contact matrices from the single-cell Hi-C data at the 64-cell stage, the single-cell HiC-Pro *validPairs* files were downloaded from GSE129029 and combined into one file, from which each valid pair was then assigned to either maternal or paternal group depending on the HiC-Pro assigned allele status. For example, if a valid pair had both reads assigned to the paternal genome or one read of the pair was mapped to the paternal and the other one was unassigned, it would be considered as a paternal valid pair. Similarly, the maternal-specific valid pairs were also identified. Once the paternal and maternal specific bulk *validPairs* files were prepared, they were converted into contact matrices (.hic format) using hicpro2juicebox utility for visualization and further downstream analysis^71^. For allele-specific compartment analysis, we extracted the Knight-Ruiz (KR) normalized contact matrices at 250 kb resolution from specific .hic files and compartment calling was performed using matrix2compartment.pl script from the cworld-dekker tools (https://github.com/dekkerlab/cworld-dekker). Similarly, for the boundary insulation score calculation from the KR normalized matrices at 50 kb resolution, matrix2insulation.pl script was used from the cworld-dekker tools.

### Xa-hyperactivation analysis

In each embryo, the TE expression on the Xa and autosomes was represented by the total counts of allelic TE reads that were assigned to the Xa and autosomes, respectively, and the TE expression from an autosome was further calculated by averaging the total autosomal TE expression. For males, all allelic reads assigned to the X were considered from the Xa. These TE expression from the Xa and an autosome was then normalized to the total autosomal gene read counts from the same embryo. To reveal the degree of TE expression change on the Xa and an autosome during imprinted XCI, the relative TE expression for the Xa and an autosome in each embryo was then calculated by further normalizing their calculated TE expression to the median value of TE expression from the Xa and an autosome at late2C stage, respectively. Similar analysis was performed for random XCI in differentiating ES cells, except that the relative expression was calculated by normalizing to the median value at Day 0 of differentiation.

### RNA FISH for LINEs

F1 embryos were harvested at expected stage from the cross between male CAST/EiJ and female C57BL/6J, followed by the removal of zona pellucida in acid Tyrode’s solution and two washes in PBS-BSA (1mg/ml). Embryos were then transferred onto defrosted glass slides with minimal carry-over liquid. Once completely dried out, embryos were then fixed and permeabilized by first incubating in 50ml of fresh 1% PFA in PBS with 0.05% Tergitol (Sigma) for 5 mins on ice, followed by transferring into 50ml of 1% PFA in PBS for another 5 min on ice. After incubation, slides were transferred to 70% ethanol on ice, and stored at 4C until use.

### Quantification and Statistical Analysis

Statistical analyses were performed by R. Statistical details and results of experiments can be found in the figure legends and figures. p < 0.05 in most statistical tests was considered significance.

