## Supplemental Figures for "Dosage compensation of transposable elements in mammals"

### SUPPLEMENTARY FIGURE LEGENDS

**Supplementary Figure 1. Sensitive detection of TE expression using so-smart-seq reveals sample reproducibility. Related to Figure 1.**

a. Comparing coverage profiles of representative TE (MERVL, B2) RNAs using our So-Smart-seq method versus a published method (SMART-seq[25]). Embryos were all at 4C stage. The best embryo replicate from each method was used for comparison.

b. Metagene analysis showing the coverage of SINEs and LTRs on chr1 using So-smart-seq (current method) and Smart-seq2[21]. Embryos were all at 8C stage. The best embryo replicate from each method were used for comparison.

c. Pairwise Pearson correlation using all expressed TEs followed by hierarchical clustering of all WT embryos (both CM and MC crosses) from So-Smart-seq. Clustering analysis is based on the Ward's method. The developmental stage and cross type of each replicate were labeled on the X and Y axis, respectively.

d. PCA analysis of differentiating ES cells at different stages.

**Supplementary Figure 2. Allelic pipeline validation and dosage compensation of TEs in CM and MC embryos. Related to Figure 1.**

a. Percentage of TE RNAs in the total transcriptome of CM preimplantation embryos. Total reads mapping to TEs (LINEs, SINEs, LTR and DNA) were compared with the total genome mapping reads (excluding rRNA reads).

b. Percentage of each TE class in the total expressed TE RNAs during preimplantation development (CM cross).

c. Percentage of TE RNAs in the total transcriptome of differentiating ES cells. Total reads mapping to TEs (LINEs, SINEs, LTR and DNA) were compared with the total genome mapping reads (excluding rRNA reads).

d. Percentage of each TE class in the total expressed TE RNAs during ES cell differentiation.

e. The distribution of allelic TE reads on the X chromosome. Each vertical blue line on the tracks represents a mapped allelic TE read.

f. Ratio of Xp versus Xm TE reads in male and female embryos across preimplantation development.

g. Determined allelic read origin in samples from pure parental strains (*mus* or *cast*) by our bioinformatic pipeline.

h. The silencing dynamics of Xp TEs in MC crossed embryos. total autosomal TEs are used as comparators. **p*<0.0005, ***p*<<0.0001, by Mann-Whitney U test.

i. The silencing dynamics of Xp TEs in CM embryos. Two autosomes (chr1 and chr13) are used as autosome comparators. **p*<0.006, ***p*<<0.001, ****p*=0.013, by Mann-Whitney U test.

j. Density plots showing the skewing of X-linked gene elements in MC and CM embryos. Gene skewing on chr13 is also included as an internal control. P values were determined by Mann-Whitney U test.

**Supplementary Figure 3. SINE and LTR expression in preimplantation embryos. Related to Figure 2 and Figure 3.**

a. Percentage of 4 TE classes in the expressed TE RNAs on the Xm (top) and Xp (bottom) through preimplantation development.

b. Heatmap showing the paternal fraction of SINE expression on autosomes and X-chromosome at zygote stage. Embryos were from MC cross.

c. Metagene analysis showing the read mapping on the loci of SINE elements.

d. SINE expression on the Xm remains unchanged during imprinted XCI (from 4C to early blastocyst). Allelic read counts were normalized to the total expressed SINEs. The vertical red dash line marks *Xist* locus.

e. Pie chart showing the fractions of SINE families expressed on the Xp using the SINE allelic reads combined from all stages.

f. Silencing dynamics of 4 dominate classes of LTRs in preimplantation embryos.

g. Abundance of full-length LINEs (>5kb) for each LINE family in both *mus* (mm10) and *cast* (GCA_921999005.2) genome. Young (L1Md) LINEs are marked in bold letters.

h. Density plot showing the length distribution of all mapped intergenic and intronic LINEs on the Xp.

i. Silencing dynamics of intergenic and intronic LINEs in preimplantation embryos, revealed by our So-smart-seq.

**Supplementary Figure 4. TE silencing dynamics in different genetic background. Related to Figure 4.**

a. Silencing dynamics of Xp LINEs in both MC and CM wildtype preimplantation embryos.

b. Silencing dynamics of Xp LTRs in both MC and CM wildtype preimplantation embryos.

c. Silencing dynamics of Xp SINEs in paternal *Xist* knockout and CM wildtype preimplantation embryos. *p* values were calculated by Mann-Whitney U test.

d. Silencing dynamics of Xp LINEs in paternal *Xist* knockout and CM wildtype preimplantation embryos. $P>0.05$ at all stages, by Mann-Whitney U test.

e. Silencing dynamics of Xp LTRs in paternal *Xist* knockout and CM wildtype preimplantation embryos. *p* values were calculated by Mann-Whitney U test.

f. LTR expression across the entire Xp (*mus* allele) in CM wildtype preimplantation embryos (from 4C to early blastocyst). X chromosome was stratified into "Proximal", "Distal_1", and "Distal_2" by the proximity to *Xist* locus. Regions were labeled with different colors. Allelic read counts were normalized to the total expressed LTRs. The vertical red dash line marks *Xist* locus.

g. Violin plots showing the Xp silencing of SINEs in different regions from 4C to 8C. *p* values were calculated by Mann-Whitney U test.

h. Violin plots showing the Xp silencing of SINEs in different regions from 8C to 16C. *p* values were calculated by Mann-Whitney U test.

**Supplementary Figure 5. Progressive TE silencing during random XCI in differentiating ES cells. Related to Figure 5.**

a. SINE expression across the entire Xi (*mus* allele) in differentiating ES cells (from Day0 to Day10). The vertical red dash line marks *Xist* locus. Escapee SINEs at Day10 and their overlapping genes are labeled. Escapee genes are marked in red.

b. Progressive silencing of SINEs in different families.

c. Silencing dynamics of gene elements that overlap with escapee TEs in differentiating ES cells. Escapee and Xi-biased genes were labeled in red.

d. LINE expression across the entire Xi (*mus* allele) in differentiating ES cells (from Day0 to Day10). The vertical red dash line marks *Xist* locus. Escapee LINEs at Day10 and their overlapping genes are labeled. Escapee genes are marked in red.

e. LTR expression across the entire Xi (*mus* allele) in differentiating ES cells (from Day0 to Day10). The vertical red dash line marks *Xist* locus. Escapee LTRs at Day10 and their overlapping genes are labeled. Escapee genes are marked in red.

f. Boxplots showing the silencing dynamics of 4 major families of LTRs during ES cell differentiation (from Day0 to Day10).

**Supplementary Figure 6. Validation of the method for determining Xa hyperactivation in TEs. Related to Figure 6.**

a. The gene expression change (as represented by the relative expression comparing to late2C stage) on Xm (Xa) versus autosomes during female embryo preimplantation development. *$p$=0.019, **$p$=0.017, ***$p$=0.016, by paired t-test. *n.s.*, not significant.

b. The gene expression change (as represented by the relative expression comparing to Day0) on Xa versus autosomes during female ES cell differentiation. *$p$<0.001, **$p$<0.01, ***$p$<0.03, by paired t-test. *n.s.*, not significant.

c. The gene expression change (as represented by the relative expression comparing to Day0) on chr13 versus autosomes during female ES cell differentiation. *n.s.*, not significant, by paired t-test.

d. The TE expression change (as represented by the relative expression comparing to late2C stage) on chr1 versus autosomes during female preimplantation development. *n.s.*, not significant, by paired t-test. Note although p value was slightly smaller than 0.05 at early blastocyst stage, Chr1 in fact showed lower expression than autosomes, consistent with the absence of hyperactivation.

e. The TE expression change (as represented by the relative expression comparing to Day0) on chr13 versus autosomes during female ES cell differentiation. P values were 0.171, 0.144 and 0.083 at Day3, Day4 and Day6, respectively. *n.s.*, not significant, by paired t-test.

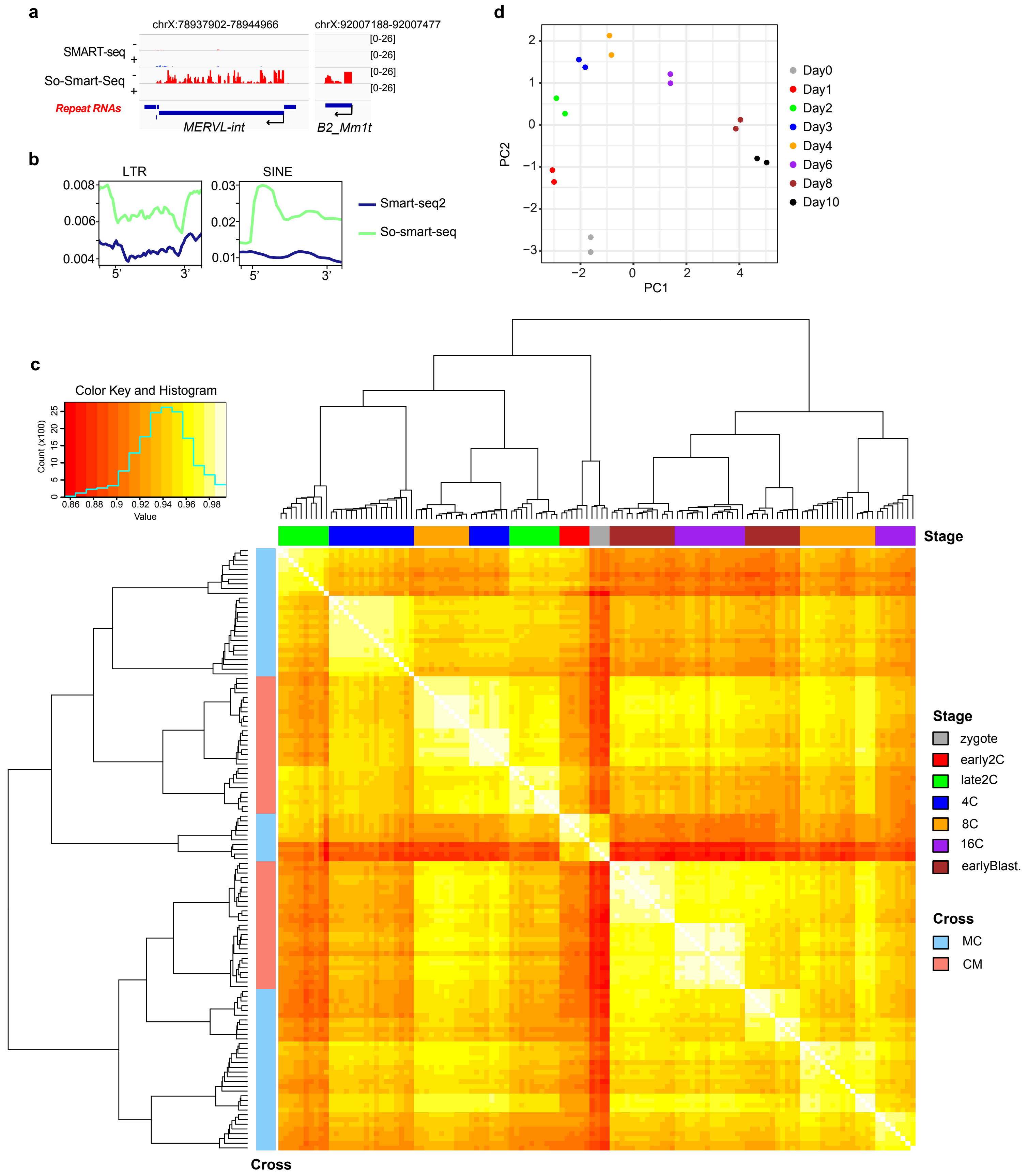


**Supplementary Fig. 1**

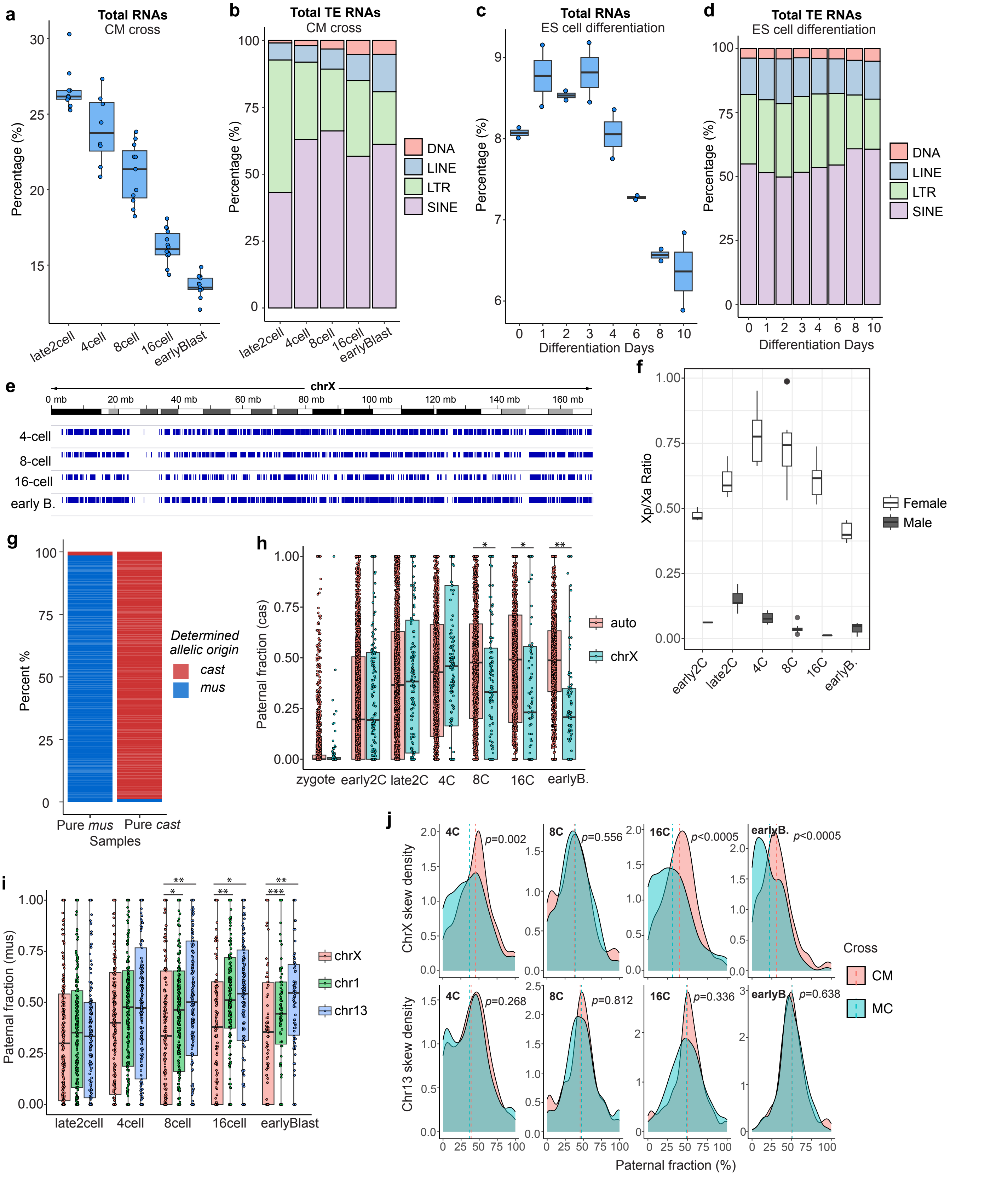


**Supplementary Fig. 2**

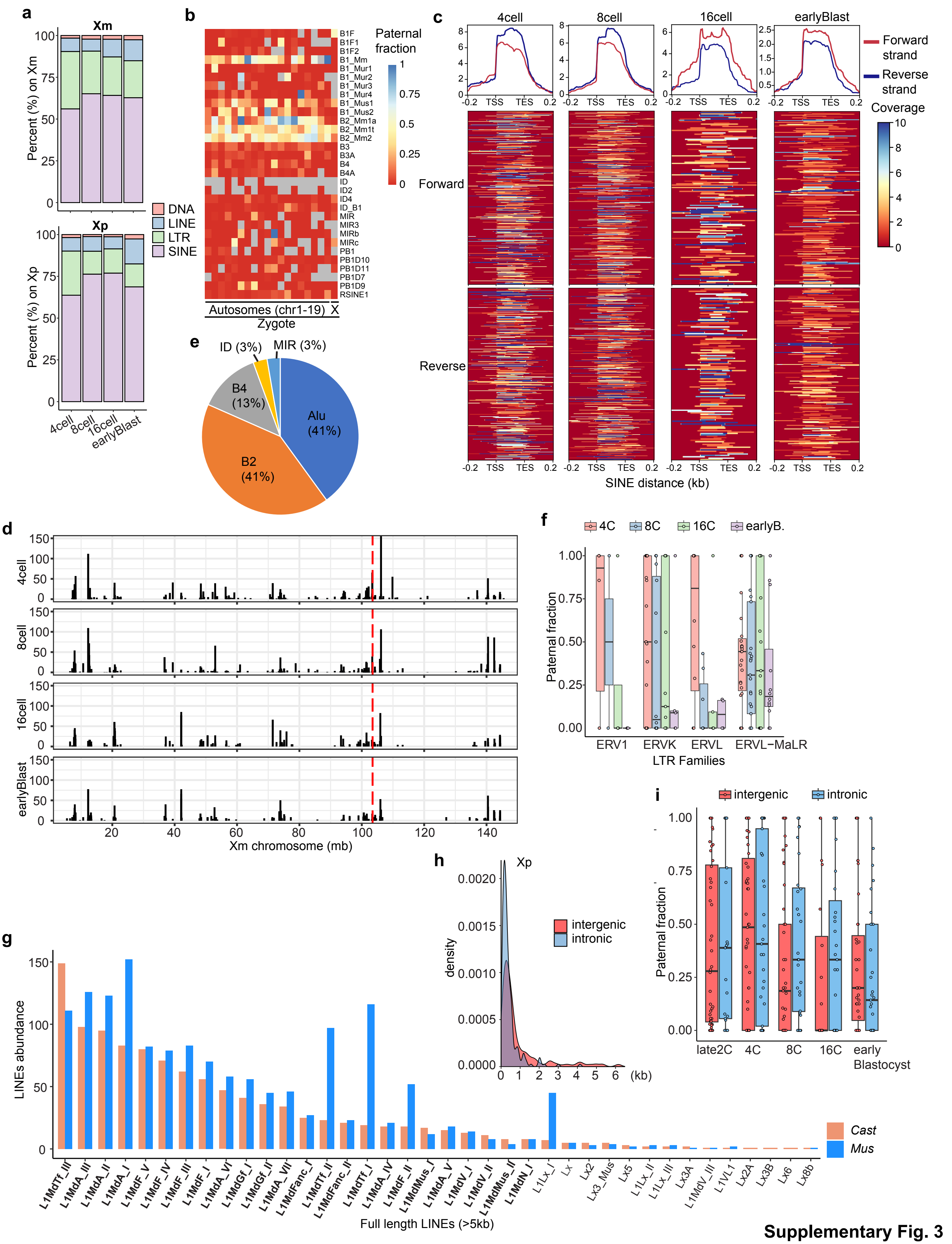


**Supplementary Fig. 3**

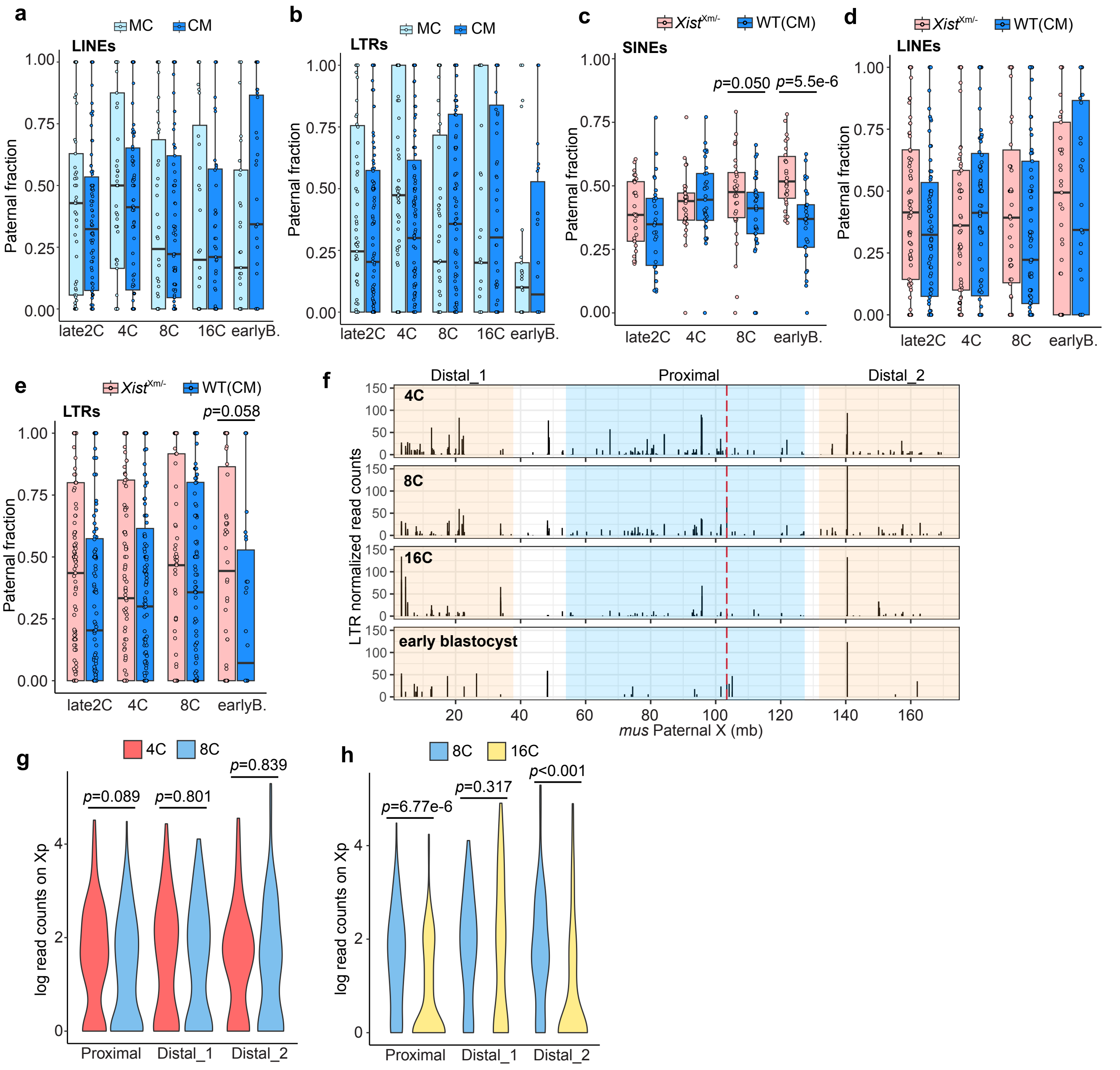


**Supplementary Fig. 4**

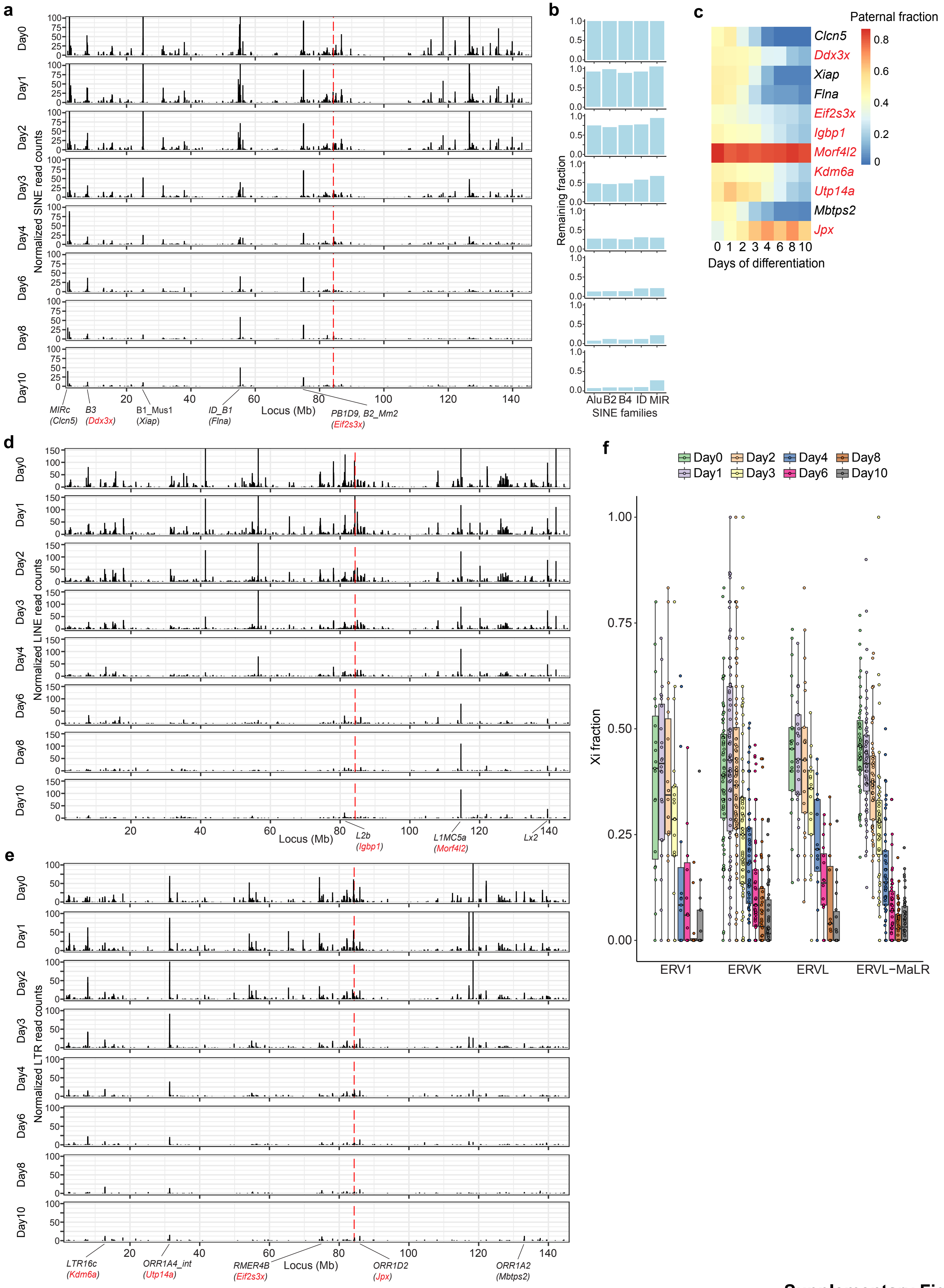


**Supplementary Fig. 5**

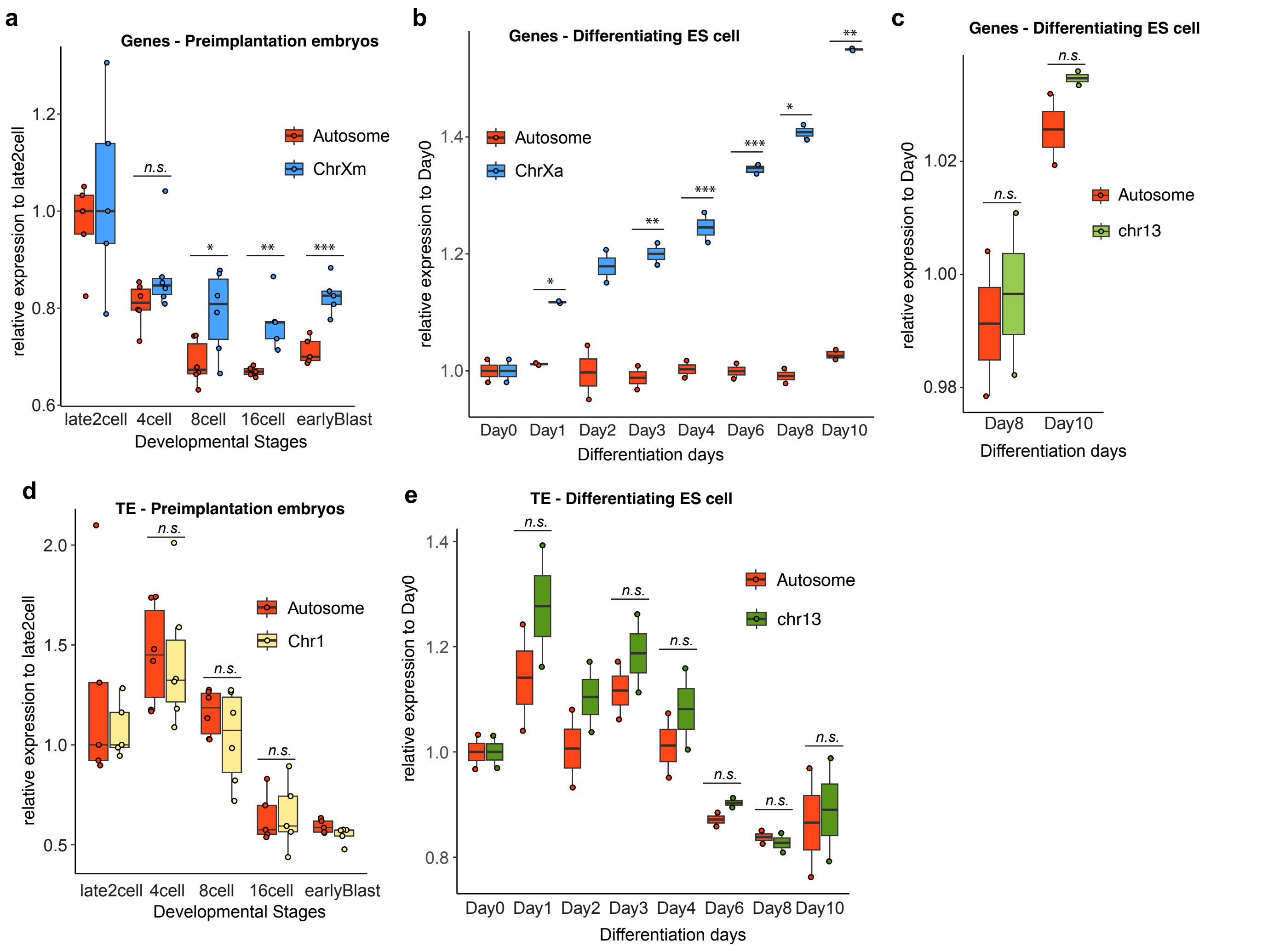


**Supplementary Fig. 6**
